# genesorteR: Feature Ranking in Clustered Single Cell Data

**DOI:** 10.1101/676379

**Authors:** Mahmoud M Ibrahim, Rafael Kramann

## Abstract

Marker genes identified in single cell experiments are expected to be highly specific to a certain cell type and highly expressed in that cell type. Detecting a gene by differential expression analysis does not necessarily satisfy those two conditions and is typically computationally expensive for large cell numbers.

Here we present genesorteR, an R package that ranks features in single cell data in a manner consistent with the expected definition of marker genes in experimental biology research. We benchmark genesorteR using various data sets and show that it is distinctly more accurate in large single cell data sets compared to other methods. genesorteR is orders of magnitude faster than current implementations of differential expression analysis methods, can operate on data containing millions of cells and is applicable to both single cell RNA-Seq and single cell ATAC-Seq data.

genesorteR is available at https://github.com/mahmoudibrahim/genesorteR.

## 1 Introduction

A main task in single cell gene expression data analysis (scRNA-Seq) is selecting marker genes after cell types are identified via unsupervised clustering of single cells. This is usually accomplished by differential expression analysis [1, 2]. On the experimental side, detecting a cell type’s marker gene by immunostaining or in a FACS experiment is often used as a proxy for identifying all cells comprising that cell type. Therefore, marker genes are expected to be highly expressed in a specific cell type and only in that cell type (ie. highly expressed and highly exclusive). Detecting a gene by differential expression analysis does not necessarily satisfy those two conditions.

Typically only a small fraction of expressed genes is captured in any single cell. The detection rate of a gene, defined as the proportion of cells in which a gene is detected, relates to its expression level (Supplementary Figures S1-S2) [3], and can be used to identify highly variable genes [4]. We propose that empirical statistics combining gene detection rate with cell clustering information can also be used to reliably estimate the specificity of genes in cell clusters.

Here we present genesorteR, an R package that provides cell type specificity gene ranking metrics that are directly related to the expected “marker gene” definition in biology. genesorteR is applicable to both scRNA-Seq and single cell ATAC-Seq (scATAC-Seq) data, can analyze thousands (millions) of cells in a few seconds (minutes) and provides various methods to select sets of marker genes and to assess cell clustering quality. genesorteR has similar accuracy to other methods in data sets containing a small number of cells, but is more accurate on large data sets consisting of thousands of cells.

## 2 Approach

### 2.1 Feature Ranking by Specificity Scores

We start with a sparse *m × n* matrix of gene expression data for *m* genes and *n* cells and a partitioning of *n* cells into *k* types *t* ∈ {*t*_1_*,…, t*_*k*_} where *k >* 1 (Figure 1a). The posterior probability of observing a cell type *t*_*i*_ given that a gene *g*_*j*_ is detected is

**Figure 1:**
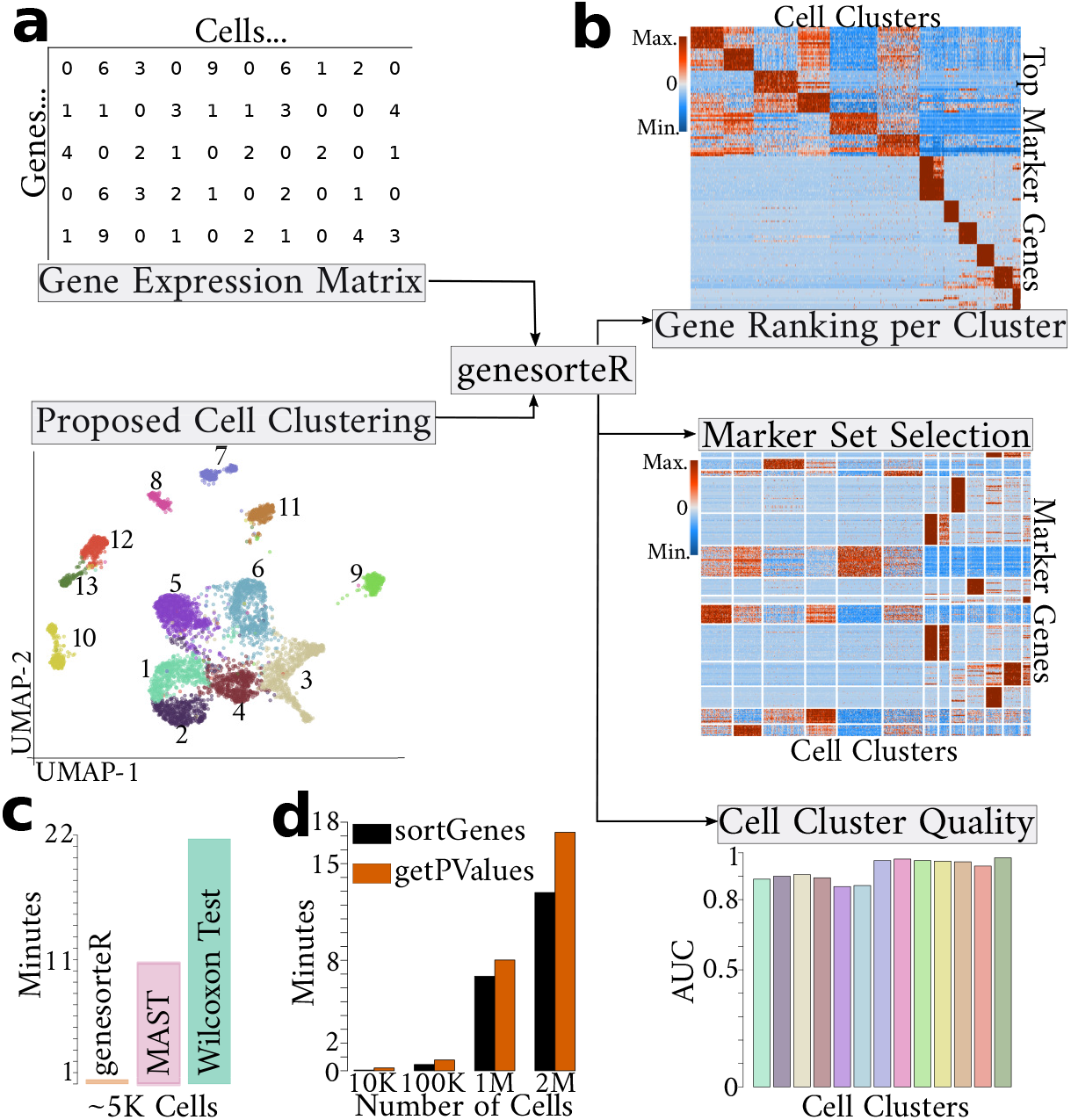
[A] The input to genesorteR is a gene expression matrix and a corresponding vector of cluster labels. [B] genesorteR, demonstrated here using 10X embryonic mouse heart data [5], can output accurate ranking of features for each cluster (top), select small or large sets of marker genes for each cell cluster (middle) and provide metrics for clustering quality (bottom). [C] Running time of genesorteR (running sortGenes+getPValues sequentially) compared to MAST [6] and Wilcoxon Rank-sum test [7] as implemented in the Seurat R package findAllMarkers function [8] (same data as [B]). [D] Running time of two main genesorteR functions on large data sets.

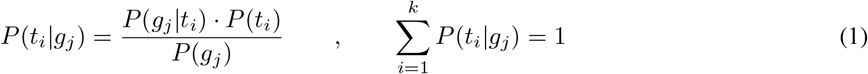

where *P*(*g*_*j*_|*t*_*i*_) is the fraction of cells assigned to *t*_*i*_ in which *g*_*j*_ was detected, *P*(*t*_*i*_) is the fraction of all *n* cells that were assigned to the partition *t*_*i*_ and *P*(*g*_*j*_) is the fraction of all *n* cells where *g*_*j*_ was detected. *P*(*t*_*i*_|*g*_*j*_) gives a measure of the exclusivity of a gene to a cell cluster: what is the probability that one has detected cell type *t*_*i*_ if gene *g*_*j*_ was detected? However, it does not address expression level. We define a Specificity Score for gene *g*_*j*_ in cell type *t*_*i*_:

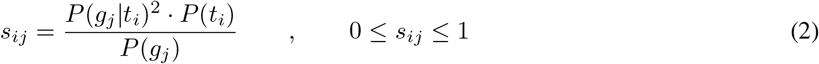

*s*_*ij*_ = 1 iff *g*_*j*_ is detected in all cells assigned to *t*_*i*_ and in none of the cells assigned to *t*_*x*≠*i*_. Therefore, *s*_*ij*_ measures whether a gene is exclusive to a cluster and whether it is highly expressed in that cluster. The genesorteR function *sortGenes* returns an *m × k* matrix *S* of specificity scores that can be used to rank all genes for each cell type by their specificity scores (Figure 1b top, Supplementary Figures S3-S4).

### 2.2 Selecting Sets of Marker Genes

A marker gene will ideally have high *s* score in only one cell type. We define an entropy-based metric for each gene *g*_*j*_ as the Gene Specificity Shannon Index:

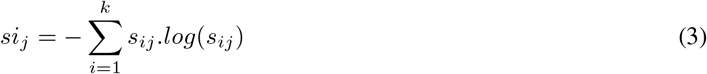

Marker genes can be selected as those with high *s* scores and low *si* scores [9]. The *getMarkers* function selects a small set of marker genes (Figure 1b, middle) by selecting the top *x* percentile of genes from *S* and partitioning them into low *si* and high *si* genes (Supplementary Figure S5, see Supplementary Text 1).

We may also be interested in an extended set of highly variable genes. The *getPValues* function performs a permutation test where the matrix *S* is recomputed after cluster labels are randomly permuted to obtain an empirical null distribution of specificity scores. *getPValues* returns an *m × k* matrix *P* of *p*-values expressing the probabilities of observing the scores in *S* if cells were randomly allocated to *k* partitions of the same sizes as the partitions obtained from clustering (Supplementary Figures S6-S7). *p*-values are calculated according to the formula for permutations without replacement provided in Phipson and Smyth [10] (see Supplementary Text 1). *getPValues* provides reasonable *p*-values with a nearly uniform distribution (Supplementary Figures S6-S7). See the Results section for detailed analysis of *getPValues* performance compared to methods implemented in the Seurat R package [8].

### 2.3 Visualization

*plotMarkerHeat* (Figure 1b, middle) makes use of gene clustering and cell averaging to quickly summarize the expression of up to tens of thousands of genes over thousands to millions of cells. *plotMarkerHeat* clusters genes using the k-means algorithm applied to the Euclidean norm-scaled gene expression vectors (see Supplementary Text 1). Similar functions for plotting binary matrices are also available.

### 2.4 Cluster Quality Evaluation

Single cell clustering results can vary widely depending on the clustering methods, parameters and the features selected for clustering. Therefore, metrics to evaluate clustering quality are highly desirable. A well-separated identifiable cluster will feature a small group of genes with relatively high specificity scores relative to all other genes in that cluster. Sorting genes by specificity scores and plotting the scaled scores in order produces a curve that should be far from the diagonal for well-separated clusters and closer to the diagonal for poor clusters (Supplementary Figure S8). The area-under-the-curve can be used to quantify this intuition (Figure 1b, bottom). The *getClassAUC* function performs this clustering quality evaluation on the output of the *sortGenes* function. This metric is inspired by metrics used to evaluate ChIP-Seq data quality in which the concentration of a large percentage of reads in a small percentage of genomic bins is an indicator of high quality data [11].

### 2.5 Implementation and Speed

genesorteR operates on sparse Matrices using the Matrix R package [12], and can process data containing thousands to millions of cells in a few seconds to a few minutes respectively (Figure 1c-d). genesorteR is orders of magnitude faster than popular single cell differential expression analysis methods (Figure 1c-d).

## 3 Results

### 3.1 Binarization Methods Influence genesorteR’s Results

It was pointed out in *Finak et al.* [6] that naive binarization of scRNA-Seq data (keeping all non-zero entries) achieves lower accuracy than binarization methods which take gene expression level into account. genesorteR implements three binarization methods within the *sortGenes* function: (1) naive binarization, (2) median binarization where the median of all non-zero entries is taken as a cutoff and (3) adaptive median binarization (inspired by MAST [6]) where genes are first clustered into distinct groups based on their expression levels and the median binarization method is applied to each group separately (see Supplementary Text 1).

To test these binarization methods, we obtained three different scRNA-Seq data sets consisting of mixes of cell types for which bulk gene expression data were also available. We used differential expression results on the bulk gene expression data as ground truth for benchmarking scRNA-Seq methods (see Methods, Supplementary Figure S9). Data from *Velasco et al.* [13] included 143 single cells from mouse embryonic bodies and induced spinal motor neurons, data from *Islam et al.* [14] included 92 cells from mouse embryonic stem cells and mouse embryonic fibroblasts and data from *Zheng et al.* [15] included 3388 cells from human Jurkat cells and human HEK293T cells (see Methods, Supplementary Figure S9).

Consistent with the observations in *Finak et al.* [6], we noticed that genesorteR’s naive binarization achieves lower accuracy than median and adaptive median binarization methods in both the *Velasco et al.* and the *Islam et al.* data (Figure 2). However, this difference in performance is almost entirely negated in the larger *Zheng et al.* data set (Figure 2). In large high-quality data that is sequenced to near saturation, naive binarization may be sufficient to achieve comparable accuracy to more sophisticated binarization methods. As expected naive binarization also consistently achieves higher sensitivity than adaptive median and median binarization methods (Supplementary Figure S10). We also note that all three methods achieve similar average log fold change ratio (Figure 3a), but the median method consistently identifies genes captured in a higher fraction of cells than naive and adaptive median methods (Figure 3b).

**Figure 2:**
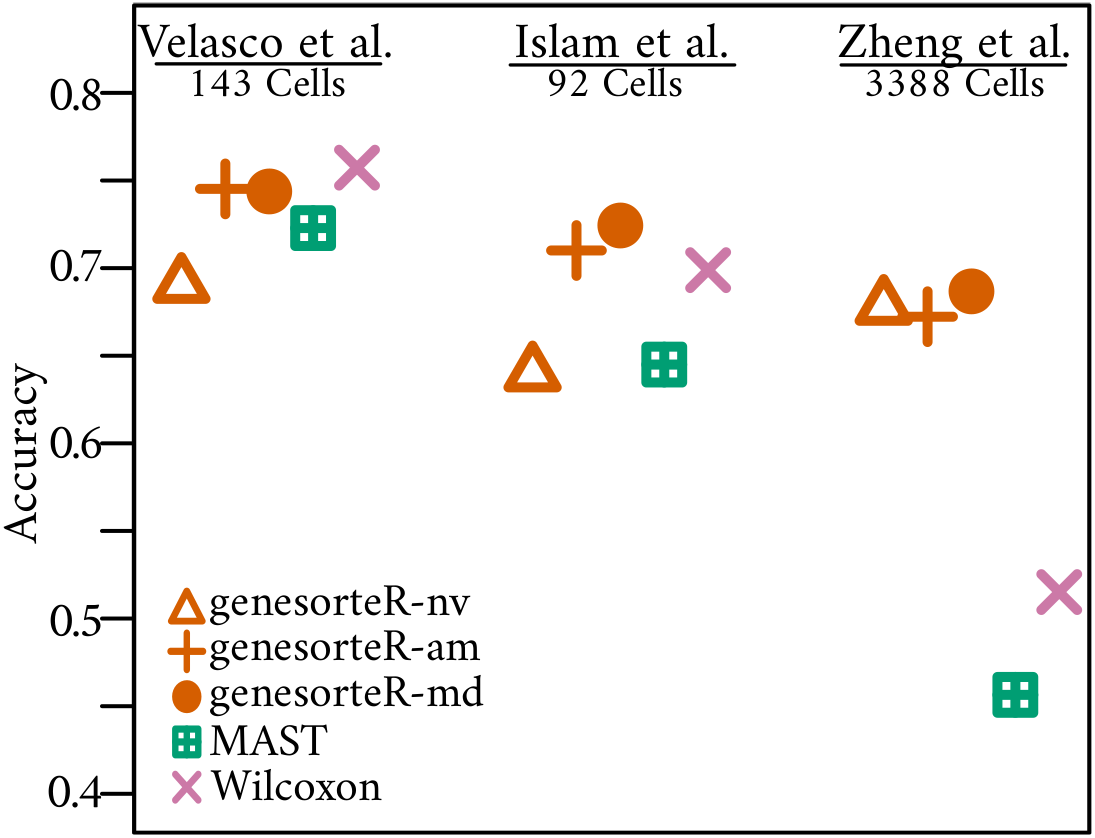
Accuracy of scRNA-Seq Differential Expression methods on various data sets. nv=naive binarization, am=adaptive median binarization, md=median binarization. MAST [6] and Wilcoxon test [7] were run through the Seurat R package [8]. Also see Supplementary Figure S10.

**Figure 3:**
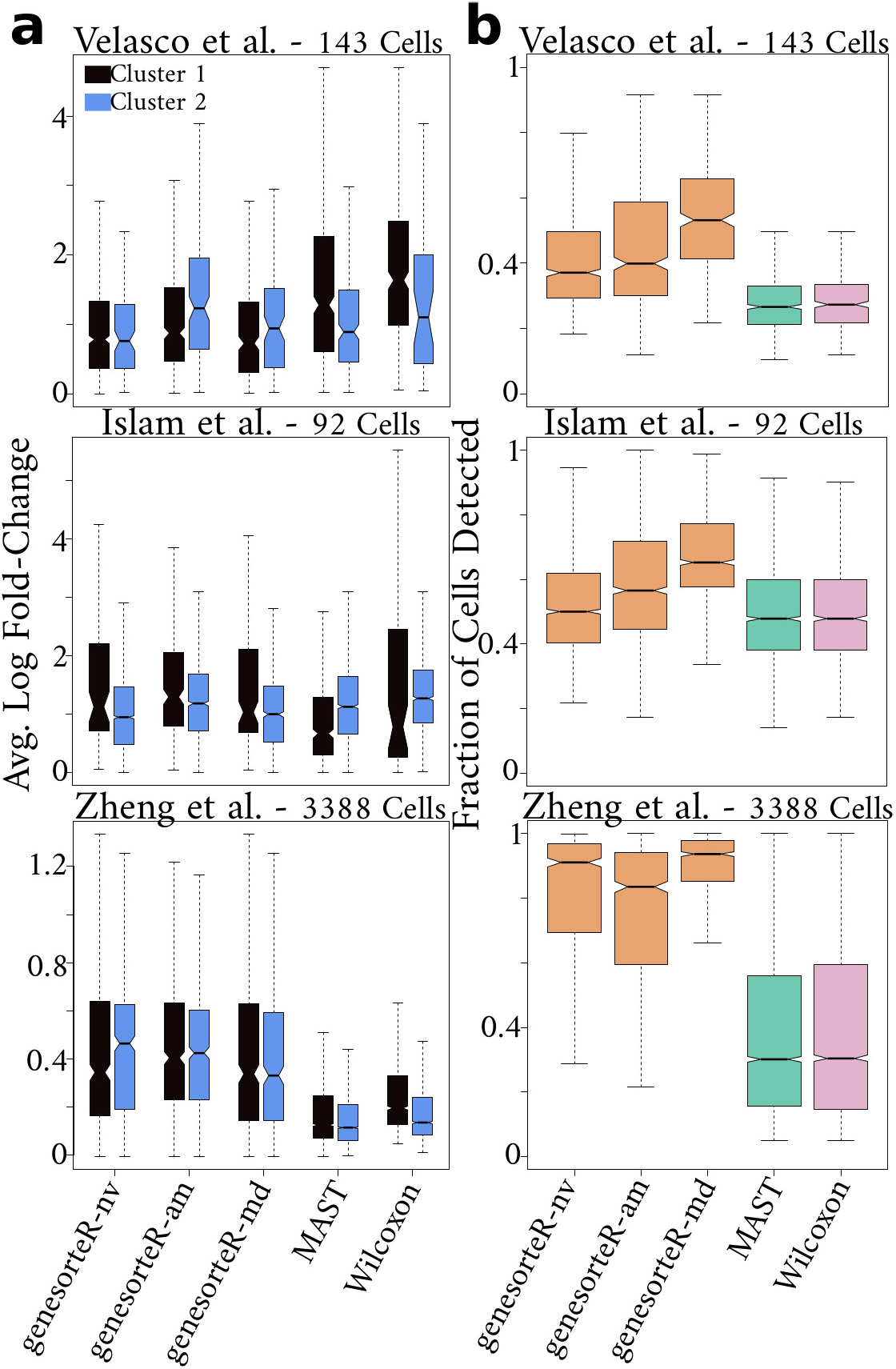
[A] Average log-fold change of differentially expressed genes. Each data set consists of two known cell clusters, highlighted in blue and black. [B] Fraction of cells in which differentially expressed genes were detected. nv=naive binarization, am=adaptive median binarization, md=median binarization. MAST [6] and Wilcoxon test [7] were run via the Seurat R package [8]. See also Supplementary Figure S11.

Overall, these analyses indicate that adaptive median and median binarization methods are more suitable for data sets with low cell numbers. With larger data sets, naive binarization will also perform comparably but the median binarization method identifies genes with higher detection rate. Median binarization is genesorteR’s default binarization method.

### 3.2 genesorteR Provides Accurate Selection of Variable Genes

Next we sought to gauge genesorteR’s performance relative to single cell differential expression analysis methods implemented in the popular Seurat R package [8], including the Wilcoxon test [7] and MAST [6]. We chose the Wilcoxon test since it is a non-parametric method and the default of the Seurat R package *findAllMarkers* function. We chose MAST since it makes use of both binary gene detection information (like genesorteR) and of gene transcript count information.

All three methods achieve comparable accuracy in smaller data sets (*Velasco et al.* [13] and *Islam et al.* [14]), but genesorteR achieves better accuracy in the larger *Zheng et al.* data set [15] (Figure 2). While MAST and the Wilcoxon test are four to five times more sensitive than genesorteR in large data, they have markedly lower specificity and precision (Supplementary Figure S10). Indeed it was recently noted that current methods for differential expression analysis on large scRNA-Seq data yield overly optimistic and artificially low *p*-values [16]. In contrast, genesorteR’s *getPValues* does not test for differences in mean expression between clusters, but effectively tests the clustering itself, yielding reasonably conservative *p*-values (Supplementary Figure S6). In large data consisting of thousands of cells, genes selected by *getPValues* are typically a subset, or near subset, of genes selected by MAST and the Wilcoxon test (Supplementary Figure S7).

### 3.3 genesorteR Enriches for Genes with Higher Detection Rate

In addition to identifying variable genes accurately, it is desirable to identify genes whose expression has higher average log fold-change values. We compared the average log fold-change values obtained from genes identified by the Wilcoxon test [7] and MAST [6] versus those identified by genesorteR. While genesorteR, MAST and Wilcoxon achieve similar average log-fold change values in smaller data sets (Figure 3a), genesorteR’s selected genes in the larger *Zheng et al.* data set [15] have higher average log-fold change values compared to genes identified by MAST and Wilcoxon test (Figure 3a). This was also the case for the 10X mouse embryonic heart data [5] (Supplementary Figure 7).

Note that in Figure 3, we calculate average log-fold change as *log*(*µ*_*c*_) − *log*(*µ*_*o*_) where *µ*_*c*_ is the average expression in the cluster in which the gene is statistically significantly higher expressed and *µ*_*o*_ is the average expression in all other cells. This formula is similar to, or exactly the same as, the formula used in the Seurat R package[8, 17]. On the other hand, in its *getTable* function, genesorteR reports average log fold change values according to *µ*_*log*_(*e*_*c*_ +1) − *µ*_*log*_(*e*_*o*_ +1), where *µ*_*log*_ denotes the average of the log values and *e*_*c*_ is the expression in the cells belonging to the cluster in which the gene is statistically significantly higher expressed and *e*_*o*_ is the expression in all other cells. The difference is that in the first formula expression values are averaged while in the second formula, expression values are averaged in log space. genesorteR’s formula is similar to the one used in the limma R package [18, 19]. We propose that this second formula is more appropriate for single cell data since it is less sensitive to outliers and more conservative, akin to the difference between the arithmetic mean (first formula) and the geometric mean (second formula). Supplementary Figure S11 provides average log fold change values calculated according to the second formula for the same genes plotted in Figure 3a.

Next we quantified the expression level of genes detected by the three methods and found that genes detected by genesorteR are consistently higher expressed in the sense that they are detected, on average, in a higher fraction of cells than genes detected by the Wilcoxon test and MAST. This is expected because genesorteR’s specificity scores explicitly incorporate gene detection rate as a key feature for gene identification while Wilcoxon test and MAST identify, primarily, differences in mean expression.

Overall, genesorteR has comparable performance to the popular Seurat R package [8] in data sets consisting of tens to hundreds of cells. In large data consisting of thousands of cells, differential gene expression methods tested here detect a large set of genes that might not be justified. For example, MAST and the Wilcoxon test together detected ~75% of genes in the 10X mouse embryonic heart data set as differentially expressed (Supplementary Figure S7). Genes detected by genesorteR are typically a subset of those detected by Wilcoxon and/or MAST, and enrich for genes with high average log fold change (Figure 3a, Supplementary Figures S7 and S11) and higher gene detection rate (Figure 3b). Although methods implemented in Seurat’s *findAllMarkers* are distinctly more sensitive than genesorteR, genesorteR is distinctly more accurate with drastically higher specificity (Supplementary Figure S10).

### 3.4 genesorteR is Applicable to a Wide Range of scRNA-Seq Protocols

Different scRNA-Seq protocols may yield data with different properties and biases. To test whether genesorteR is applicable to a wide range of scRNA-Seq methods, we demonstrate genesorteR on various data sets including 10X chromium data (~5k cells, Figure 1a-c, Supplementary Figures S3-S8) [5], Cel-Seq2 data (~50 cells, Supplementary Figure S12) [20], Smart-Seq2 data (~45k cells, Supplementary Figure S13) [21] and SPLiT-Seq data (~100k cells, Supplementary Figure S14) [22]. This indicates that genesorteR is widely applicable to various scRNA-Seq protocols.

### 3.5 genesorteR is Applicable to scATAC-Seq Data

Since scATAC-Seq data share many of the properties of scRNA-Seq data, we wondered whether genesorteR might be applicable to scATAC-Seq data. To obtain a benchmarking data set, we generated a scATAC-Seq data set composed of an artificial mix of the two human cell lines K562 and HEK293T at an approximate ratio of 7:3 respectively (see Methods, Supplementary Figure S15). Mixing two widely characterized cell lines allows for clustering of single nuclei based on information from orthogonal public DNase-Seq data from the same cell lines. This provides ground truth for benchmarking scATAC-Seq data analysis methods.

We summarized the K562-HEK293T scATAC-Seq mix data by counting reads in 5kb windows in single nuclei and clustered the nuclei based on intersections of the detected regions in each nucleus with K562 and HEK293T specific peaks identified from DNase-Seq data (see Methods, Figure 4a). This allowed us to identify 3 groups in a semi-supervised manner (see Methods): K562 nuclei (n=3959), HEK293T nuclei (n=1883) and Ambiguous nuclei (n=344) likely arising from doublets or droplets occupied by nuclei fragments (Figure 4a,b). This data set provides a convenient example to benchmark scATAC-Seq clustering methods and, more importantly here, to benchmark marker feature detection. In the following comparisons, we measured genesorteR’s accuracy in identifying marker genomic bins against differential peak calling results on bulk DNase-Seq data (see Methods, Figure 4b).

**Figure 4:**
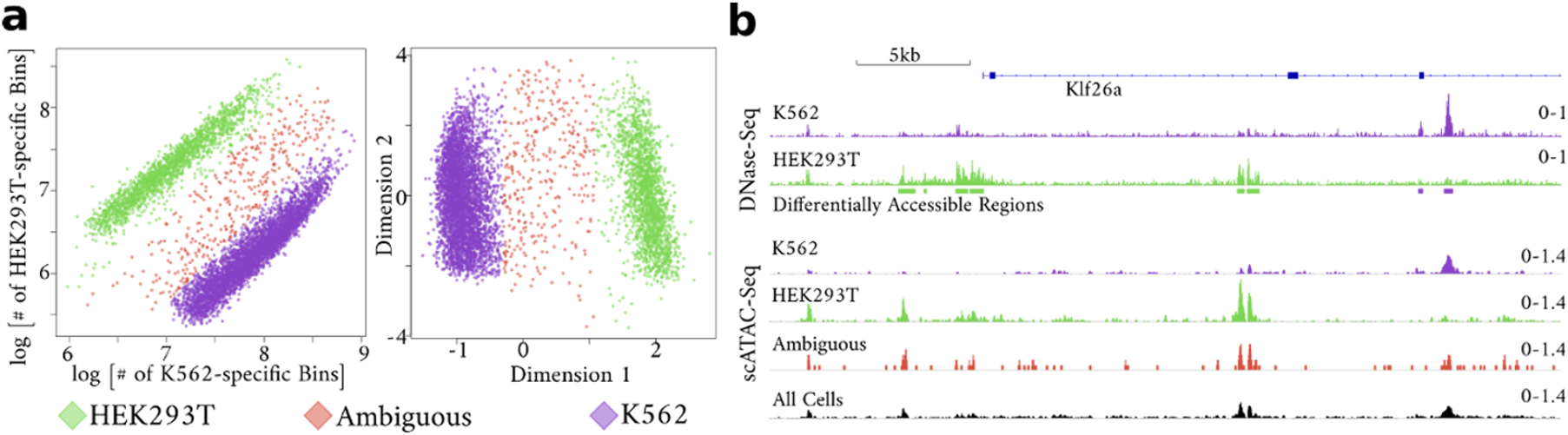
Semi-supervised clustering of scATAC-Seq data. [A] The number of genomic bins in each nucleus that intersected K562-specific and HEK293T-specific peaks obtained from DNase-Seq data [23] (left), and the left singular vectors obtained by applying Singular Value Decomposition to this peak intersection count matrix (right). Cell clusters were obtained by k-means clustering of the cells along the first singular vector. [B] An example locus showing normalized signal for K562 and HEK293T DNase-Seq data and normalized signal aggregated from nuclei belonging to the three different groups defined in scATAC-Seq data. Differentially Accessible Regions refer to differentially accessible peaks identified from DNase-Seq data.

Although segmenting the genome in equal sized bins is an intuitive way to summarize scATAC-Seq data in an unbiased manner [24], it is not immediately clear which bin size is appropriate. We observe that, as the bin size increases (and the resolution at which we summarize the data decreases), genesorteR’s accuracy decreases (Figure 5). However, a 20x increase in bin size, causes approximately only a 10% decrease in accuracy (Figure 5). Although this data set is relatively not complex with only two main cell types, this analysis indicates that as data sets become large with hundreds of thousands of nuclei, it may be justified in some cases to summarize the data at a coarse resolution for initial cell clustering analysis in order to reduce computational cost.

**Figure 5:**
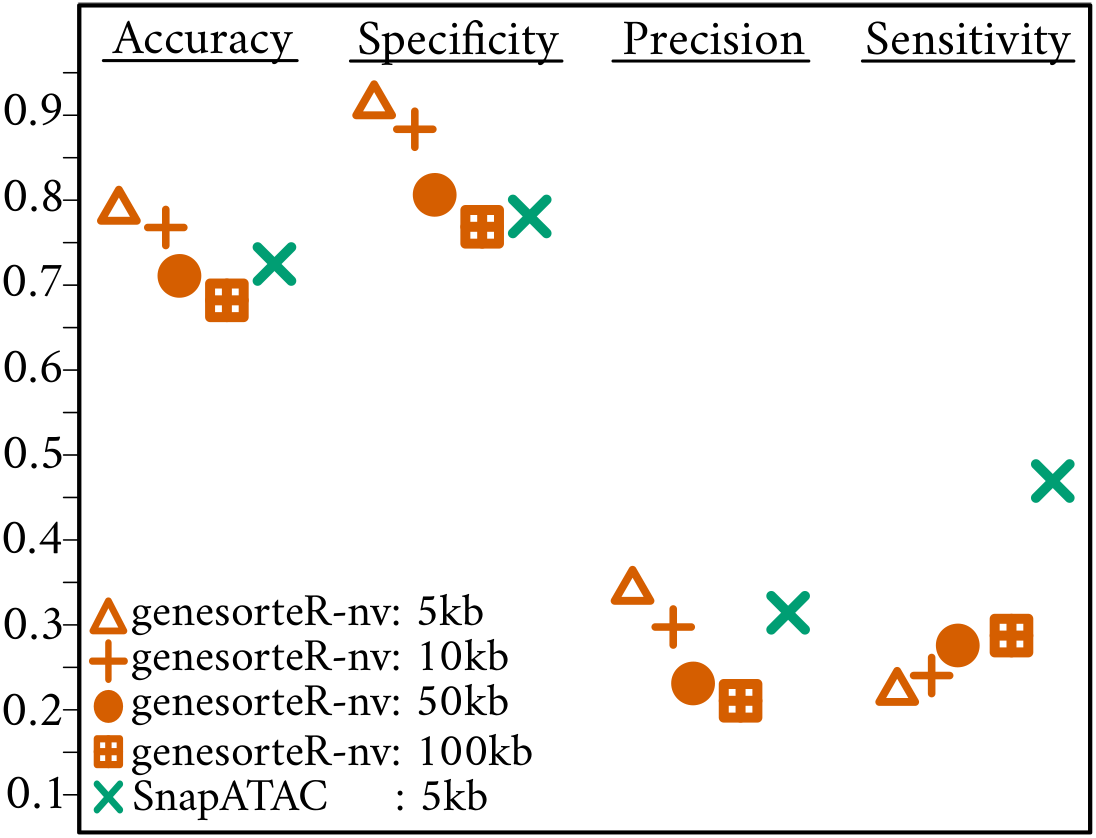
Accuracy, specificity, precision and sensitivity for defining differentially accessible regions in scATAC-Seq data at different bin sizes. Data: K562-HEK293T mix data (this paper, Ambiguous cells were not considered) benchmarked using bulk DNase-Seq data [23]. SnapATAC refers to *findDAR* function in SnapATAC R package [24].

We compared genesorteR to the recent SnapATAC scATAC-Seq analysis R package, using SnapATAC’s *findDAR* (find Differentially Accessible Regions) function [24]. SnapATAC’s accuracy at a 5kb bin size is comparable to genesorteR’s accuracy at a 50kb bin size, and genesorteR is more accurate than SnapATAC at the same bin size (Figure 5). However, as was the case with Seurat’s *findMarkers* function (see above), SnapATAC is markedly more sensitive than genesorteR although this comes at the cost of markedly decreased specificity (Figure 5).

In summary, using a high-quality benchmark scATAC-Seq data set composed of an artificial mix of two well-studied cell lines (Figure 4), we show that genesorteR is applicable to marker detection in scATAC-Seq data and that its performance is at least on par with, if not better than, state-of-the-art software designed specifically for scATAC-Seq data (Figure 5).

### 3.6 genesorteR is a General Purpose Software

Although genesorteR was developed with single cell data in mind, it is generally applicable to sparse bipartite graphs where one of the parts is clustered. Therefore, it may be used for knowledge extraction in a wide variety of contexts.

As an example, we applied genesorteR’s *sortGenes* to data from the Internet Movie Database (www.imdb.com) dating to 2005 [25]. This data is a binary sparse matrix including >220k titles and >730k actors describing which actors appear in which titles. Titles are grouped into genres. We used genesorteR to find the “top actor” for each genre. In this context, top actors are those who are “specific” to a genre; they tend to appear somewhat exclusively in titles belonging to this genre and are relatively prolific within this genre. The database also lists non-professional actors who appeared as themselves in titles such as Music films and documentaries. Paul McCartney is the top “actor” for Music titles, while the late Peter Falk who portrayed Lieutenant Columbo is the top actor for the Mystery genre (Supplementary Figure S16). By correlating genres based on actor specificity scores, we could provide a reasonable clustering of genres (Supplementary Figure S17).

## 4 Discussion

One challenge in single cell data analysis is to find differentially expressed genes despite data sparsity. Here we show that empirical statistics on gene detection rate can in fact provide accurate estimates of gene-cell type specificity. This is consistent with the recent observation that droplet scRNA-Seq data is not zero-inflated [26], indicating that expression “drop-outs” tend to be biologically driven rather than technically driven. Gene detection rate correlates with gene expression level, often following a sigmoid-like curve [3, 4], regardless of the scRNA-Seq protocol used (Supplementary Figure S1). When analyzing gene expression data, genesorteR relies on this relationship. The stronger the relationship is, the better genesorteR will perform. If drop-outs were to occur completely regardless of RNA copy number, driven purely by technical factors, genesorteR would fail to find marker genes. Indeed, we observe that genesorteR performs marginally worse than the Wilcoxon test [7, 8] in the *Velasco et al.* [13] data set, in which the relationship between read counts and gene detection rate seems to hold the least out of all the data sets we tested in this manuscript (Supplementary Figure S9).

genesorteR also relies on data sparsity. If almost all genes were detected in almost all cells, the specificity score cannot be calculated. However, we showed that further sparsity can be reasonably imposed on scRNA-Seq data using methods to threshold gene expression (see Results section 3.1). As single cell technologies develop, new protocols may result in varying data characteristics. In particular, we expect the single cell research community will move toward larger data sets with more cells. genesorteR has a distinct advantage in large data in terms of accuracy and in terms of speed; it is about ten times as fast as methods implemented in the Seurat R package [8]. This improvement in speed is because genesorteR’s specificity scores are straightforward to calculate and because of sparse matrix handling utilities in the R language. genesorteR makes heavy use of the Matrix R package for sparse matrix routines [12].

Clustering is an indispensable exploratory approach in single cell data analysis. But clustering is an ill-posed problem. Regardless of the clustering method, single cell data clustering results should always be investigated and validated carefully given the context in which the data was produced. Claims of novel cell types may not be convincing especially when not substantiated by orthogonal data and appropriate sample sizes [16, 27]. Single cell data is expensive and tedious to produce and analyze, limiting sample sizes. On the computational side, methods to evaluate cell clustering quality are needed. genesorteR provides metrics for cell clustering quality evaluation (Figure 1b, bottom, Supplementary Text 1), based on the assumption that a well-defined cell cluster can be uniquely identified using relatively few marker (“specific”) genes. Indeed, a cell type is often practically defined in experimental research by the expression of one or few marker genes that uniquely identify it. genesorteR ranks genes in a manner consistent with this marker gene definition.

In scATAC-Seq data, the feature read count is assumed to be uninformative and data is often binarized prior to analysis [28], making feature detection rate the key informative aspect of the data. genesorteR is naturally applicable to such cases. We showed that genesorteR performs better than state-of-the-art scATAC-Seq tools in determining differentially accessible regions (Figure 5). In fact we have demonstrated that genesorteR can be seen as a general purpose software for summarizing clustered sparse bipartite graphs. The definition of gene specificity we describe here (that a marker gene should be unique to a cluster, yet highly expressed in that cluster) is motivated by experimental biology research. However, it is also a general idea with similarities to text mining methods such as the TF-IDF method [29] and "Unique Elite" keywords proposed in a concurrent manuscript [30]. The genesorteR R package may indeed be useful in some contexts outside single cell research.

## 5 Conclusion

In this paper we introduced genesorteR, an R package for ranking and determining significantly variable features in clustered single cell data. genesorteR is applicable to both scRNA-Seq and scATAC-Seq data, and has significant advantages in large data sets consisting of thousands to millions of cells where it is considerably faster and more accurate than other methods. Furthermore, its output is intuitive and consistent with the expected definition of marker genes in experimental biology research.

## 6 Methods

Mouse heart data (Figure 1) and Jurkat-HEK293T data [15] were obtained as filtered gene expression matrices from the 10X Genomics website [5, 31]. Data were clustered as described in Supplementary Text 2. Seurat’s *finalAllMarkers* function version 3.0.1 [8] was used to obtain results from the Wilcoxon Rank-sum test [7] and MAST [6] setting only.pos to TRUE. For all benchmarking comparisons results were thresholded to include only genes with positive average log-fold change according to the Seurat formula [8, 17]. *Velasco et al.* data [13] was benchmarked using bulk RNA-Seq data from the same manuscript, *Islam et al.* data [14] was benchmarked using GRO-Seq data [32] and *Zheng et al.* data [15] was benchmarked using bulk RNA-Seq data obtained from the ARCHS4 database [33].

For Figure 1d, cells were sampled from SPLiT-Seq data [22] and clusters were randomly assigned as follows (cell number/cluster number): 0.1K/9 1K/18 10K/35 100K/75 1M/159 2M/177. Running time was evaluated using a single compute core.

Accuracy is (*TP* + *TN*)/(*TP* + *TN* + *FP* + *FN*), precision is *TP*/(*TP* + *FP*), specificity is *TN*/(*TN* + *FP*) and sensitivity is *TP*/(*TP* + *FN*) where *TP* is true positives, *TN* is true negatives, *FP* is false positives and *FN* is false negatives.

K562 and HEK293T mix scATAC-Seq data was produced using the 10X Genomics scATAC-Seq kit version 1.0 (10X Genomics, Product Codes 1000084, 1000086 and 1000111). K562 cells (ATCC, CCL243, Lot# 70000192) and HEK293T cells (ATCC, CRL3216, Lot# 70008735) were cultured in RPMI1650 media (Gibco, Catalog# 31870-025) and DMEM+Glutamax media (Gibco, Catalog# 31966-021) respectively, supplemented with 10% fetal bovine serum.

Raw fastq files were processed using Cell Ranger ATAC version 1.0 (10X Genomics) and fragments were counted across genomics bins from the “possorted.bam” file from Cellranger using snaptools version 1.4.7 [24]. Bulk DNase-Seq data were obtained from the ENCODE consortium UCSC webpage (http://www.genome.ucsc.edu/ENCODE/) [23] and peaks were defined using JAMM version 1.0.7rev5 [34]. BigWig files for single cell populations were produced using SnapATAC version 1.0.0 via MACS2 version 2.1.0.20151222 [35] and visualized in the Integrative Genomics Viewer (IGV) [36]. BigWig files for DNase-Seq data were produced using JAMM’s SignalGenerator script [34]. SnapATAC *findDAR* function was run setting method to “exacttest”.

All differential gene and peak expression analyses for benchmark data were performed using edgeR R package version 3.24.3 [37]. genesorteR version 0.3.3 was used for all analyses in this manuscript. R version 3.5.3 [38] was used for all analysis in this manuscript.

For further methods details, please see Supplementary Text 2.

## Acknowledgements

We would like to thank Ivan G. Costa for critical reading of the manuscript and Christoph Kuppe for help with the 10X Chromium chip and controller.

