## Supplemental Text and Figures for "genesorteR: Feature Ranking in Clustered Single Cell Data"

**Fig. S1**

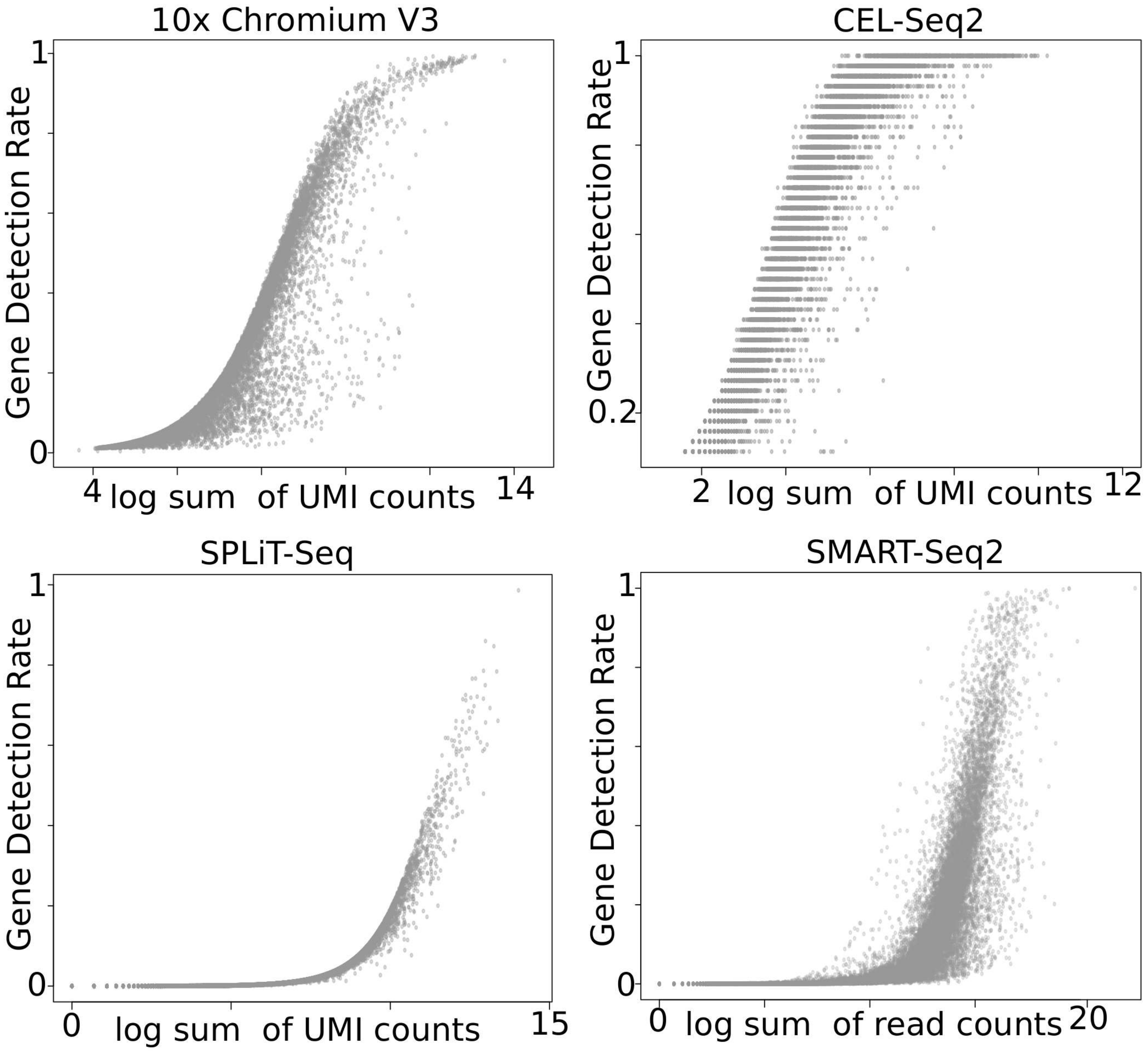
10x Chromium V3
Gene Detection Rate
1
0
4
log sum of UMI counts
14
CEL-Seq2
Gene Detection Rate
1
0.2
2
log sum of UMI counts
12
SPLiT-Seq
Gene Detection Rate
1
0
0
log sum of UMI counts
15
SMART-Seq2
Gene Detection Rate
1
0
0
log sum of read counts
20

**<u>Figure S1.</u>** Scatter plots of gene detection rate (1 – drop-out rate) versus the total number of UMI/reads detected per gene reveal that gene detection rate relates to the overall gene expression in single-cell RNA-Seq data. The exact relationship between gene detection rate and gene expression may vary based on the data generation protocol, or the number of cells or both. 10x Chromium V3 data: 5070 cells/13846 genes. SPLiT-Seq data: 95005 nuclei/26894 genes. CEL-Seq2 data: 44 cells/11420 genes. Smart-Seq2 data: 43910 cells/23341 genes. For more details on data, see Supplementary Text 2.

**Fig. S2**

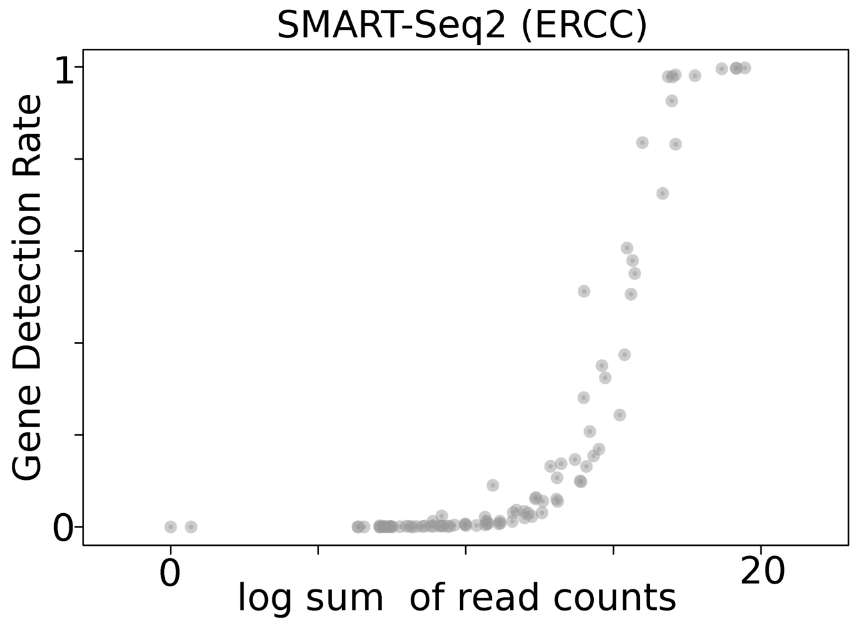
SMART-Seq2 (ERCC)
Gene Detection Rate
1
0
0
20
log sum of read counts

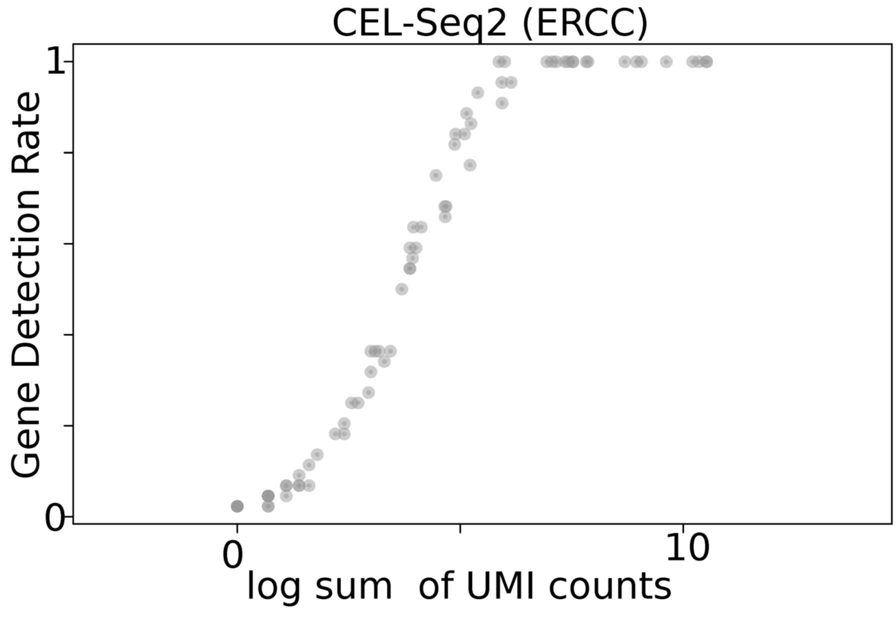
CEL-Seq2 (ERCC)
Gene Detection Rate
1
0
0
10
log sum of UMI counts

**Figure S2.** The same as Figure S1 but for ERCC transcripts associated with the CEL-Seq2 and the Smart-Seq2 datasets in Figure S1. ERCC transcripts detection rates seem to follow a sigmoidal or a near-sigmoidal relationship with their total UMI/read count.

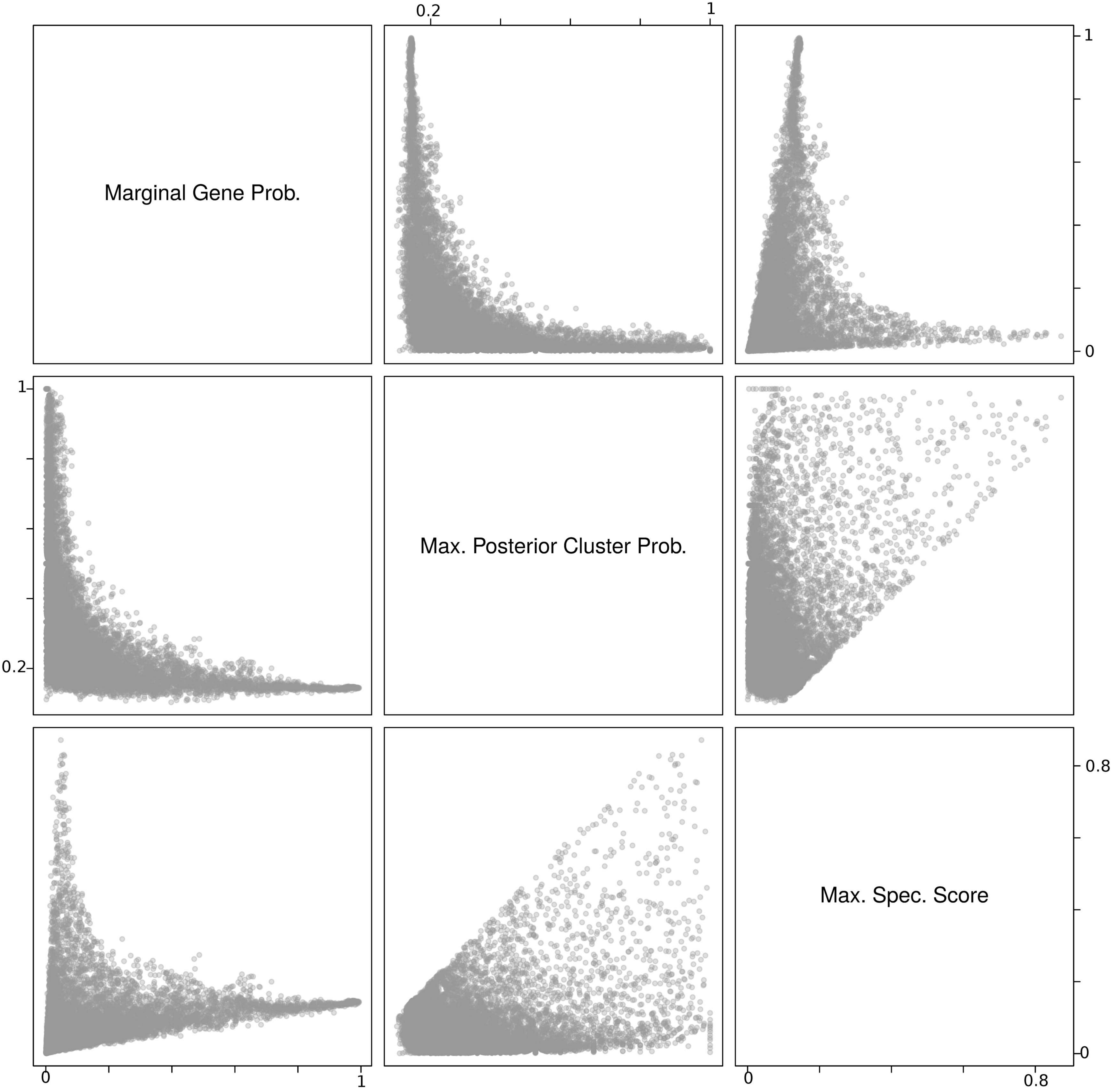
Marginal Gene Prob.
Max. Posterior Cluster Prob.
Max. Spec. Score
0.2
1
0
0.8

**<u>Figure S3.</u>** Scatter plots of the marginal gene probability (ie. gene detection rate), and the corresponding maximum posterior probability of cell clusters given the gene, and the maximum gene specificity score. Data used here is the 10x embryonic mouse heart 10k cells data, see Supplementary Text 2 for more details.

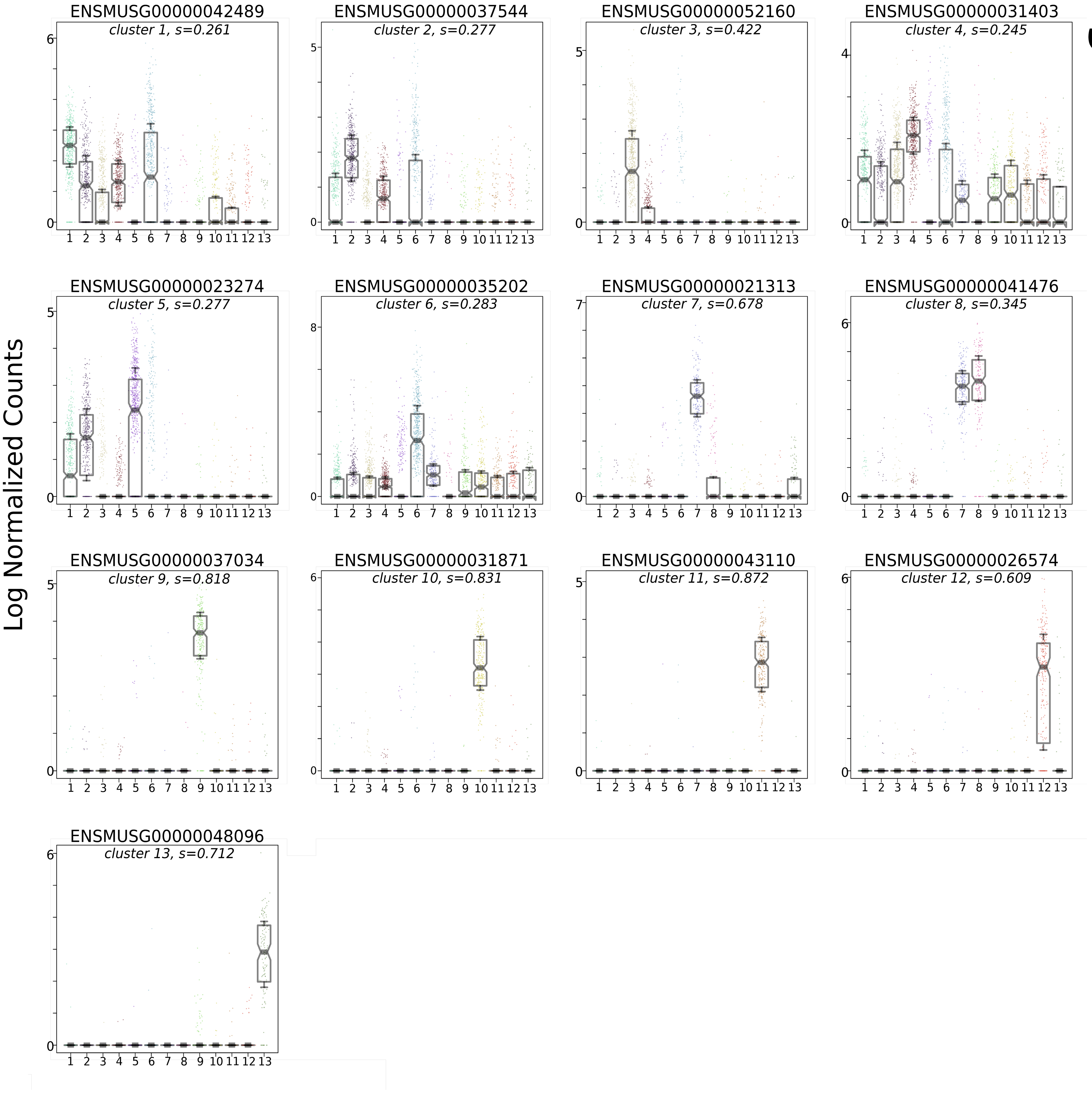
Log Normalized Counts
ENSMUSG00000042489
cluster 1, s=0.261
ENSMUSG00000037544
cluster 2, s=0.277
ENSMUSG00000052160
cluster 3, s=0.422
ENSMUSG00000031403
cluster 4, s=0.245
ENSMUSG00000023274
cluster 5, s=0.277
ENSMUSG00000035202
cluster 6, s=0.283
ENSMUSG00000021313
cluster 7, s=0.678
ENSMUSG00000041476
cluster 8, s=0.345
ENSMUSG00000037034
cluster 9, s=0.818
ENSMUSG00000031871
cluster 10, s=0.831
ENSMUSG00000043110
cluster 11, s=0.872
ENSMUSG00000026574
cluster 12, s=0.609
ENSMUSG00000048096
cluster 13, s=0.712

Fig. S4

**Figure S4.** The expression of the top marker gene in each cell cluster based on the gene specificity scores (10x 10k mouse embryonic heart data, see Supplementary Text 2). The specificity score of the gene in the corresponding cluster is indicated by “s”. The top marker genes in well-separated clusters have higher specificity scores than top marker genes in less well-separated clusters.  The specificity score accurately captures both the gene expression level in the cluster and whether it is exclusive to that cluster.

Fig. S5

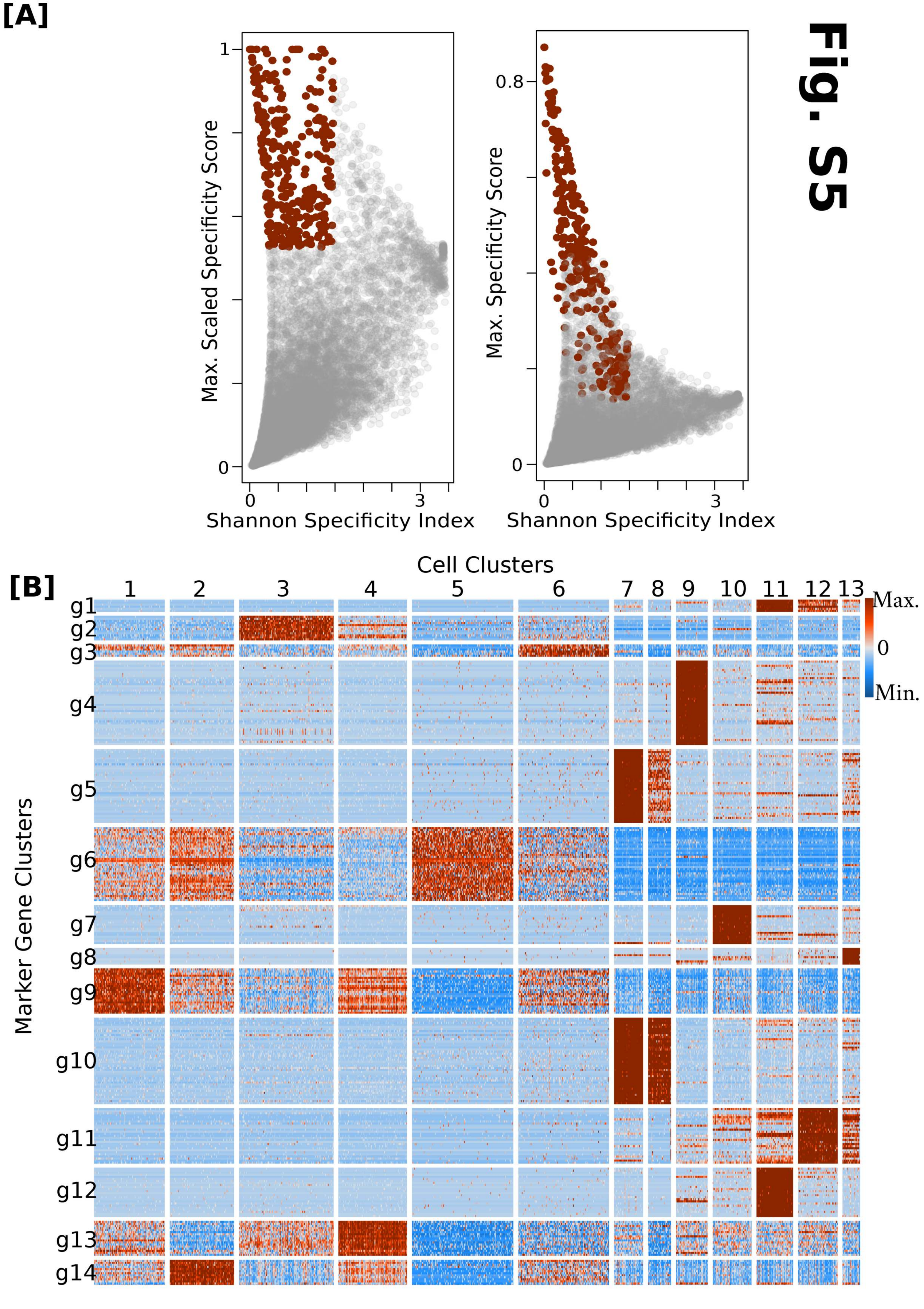

[A]
1
0
0
3
Max. Scaled Specificity Score
Shannon Specificity Index
0.8
0
0
3
Max. Specificity Score
Shannon Specificity Index
[B]
Cell Clusters
1
2
3
4
5
6
7
8
9
10
11
12
13
Marker Gene Clusters
g1
g2
g3
g4
g5
g6
g7
g8
g9
g10
g11
g12
g13
g14
Max.
0
Min.

**Figure S5.** An illustration of *getMarker()* function using 10x embryonic mouse heart data (see Supplementary Text 2). **[A]** We select the top 5% of genes based on the per-cluster scaled specificity score. Specificity scores are scaled so that the lowest score and highest score in each cell cluster is 0 and 1 respectively. The top 5% of genes are partitioned to high and low Shannon Index genes. Low Shannon index genes (red) are selected as marker genes (see Supplementary Text 1). This plot is generated using *plotMarkerScores()* function in the genesorteR package. **[B]** A heatmap of the scaled expression of genes selected using the procedure in **[A]**, genes were plotted and clustered using the *plotMarkerHeat()* function from the genesorteR package. Gene clusters in this case can be used to identify potential marker sets for experimental validation and cell population identification in FACS or immunostaining experiments. For example, genes in gene cluster “g2” are candidates to validate cell cluster 3.

Fig. S6

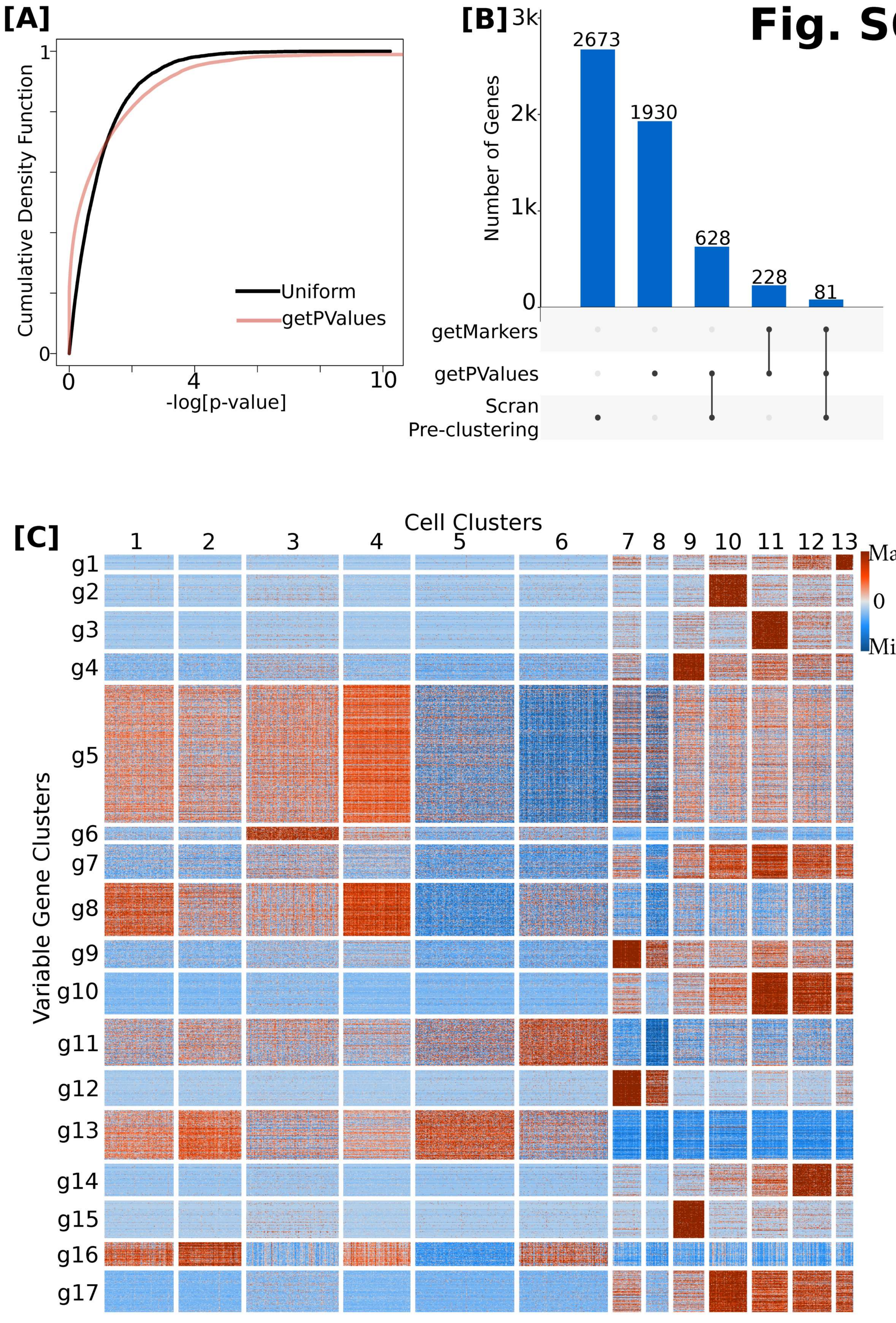

[A]
Cumulative Density Function
1
0
0
4
10
-log[p-value]
Uniform
getPValues
[B]
3k
2k
1k
0
Number of Genes
2673
1930
628
228
81
getMarkers
getPValues
Scran
Pre-clustering
[C]
Cell Clusters
1
2
3
4
5
6
7
8
9
10
11
12
13
Variable Gene Clusters
g1
g2
g3
g4
g5
g6
g7
g8
g9
g10
g11
g12
g13
g14
g15
g16
g17
Max.
0
Min.

**Figure S6.** An illustration of the *getPValues()* function from the genesorteR package on the 10x embryonic mouse heart data (see Supplementary Text 2). **[A]** *p*-values obtained using *getPValues()* are close to uniform. The abundance of conservative *p*-values is because *getPValues()* performs a one-sided test (specificity score > than what would be expected by random). **[B]** The intersection between highly variable genes identified using the Scran R package without information about clustering, the variable genes identified using genesorteR's *getPValues()* which uses clustering information and the small set of marker genes identified using genesorteR's *getMarkers()* (see **Figure S5**, see Supplementary Text 2 for more details). **[C]** A heatmap of the scaled gene expression of all genes identified by *getPValues()*, plotted and clustered using *plotMarkerHeat()*.

Fig. S7

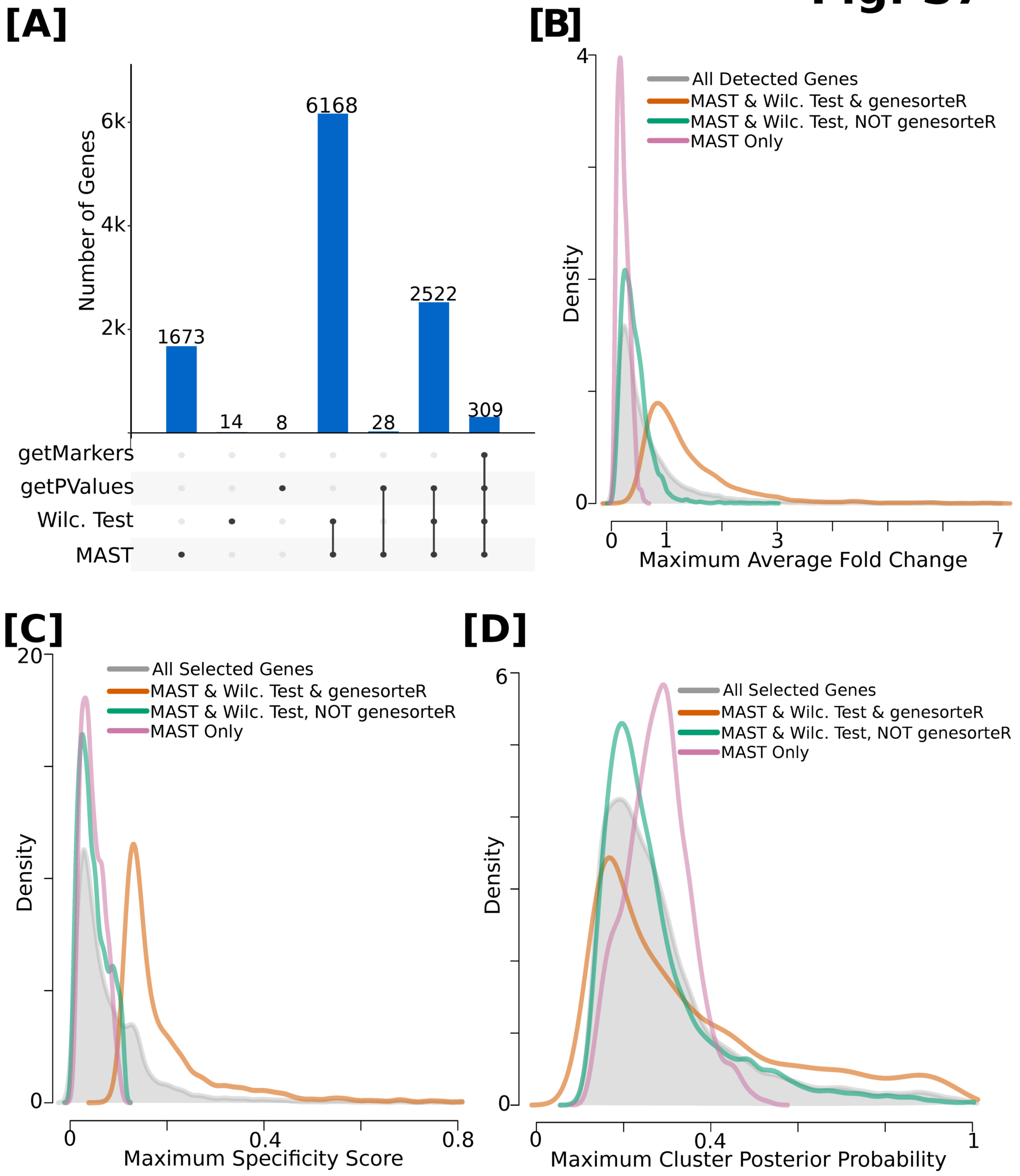

[A]
Number of Genes
6k
4k
2k
1673
14
8
6168
28
2522
309
getMarkers
getPValues
Wilc. Test
MAST
[B]
4
0
Density
All Detected Genes
MAST & Wilc. Test & genesorteR
MAST & Wilc. Test, NOT genesorteR
MAST Only
0
1
3
7
Maximum Average Fold Change
[C]
20
0
Density
All Selected Genes
MAST & Wilc. Test & genesorteR
MAST & Wilc. Test, NOT genesorteR
MAST Only
0
0.4
0.8
Maximum Specificity Score
[D]
6
0
Density
All Selected Genes
MAST & Wilc. Test & genesorteR
MAST & Wilc. Test, NOT genesorteR
MAST Only
0
0.4
1
Maximum Cluster Posterior Probability

**Figure S7.** A comparison of differential expression and marker identification methods implemented in genesorteR (getPValues and getMarkers) and Seurat (Wilcoxon test and MAST). **[A]** The intersection between genes selected by all 4 methods (at a corrected *p*-value threshold of 0.05 for getPValues, Wilcoxon test and MAST). genesorteR's *getPValues()* is more conservative such that almost all genes selected by *getPValues()* were also selected by Wilcoxon test and MAST. On the other hand, Wilcoxon test and MAST together selected ~77% of all the genes present in the data as differentially expressed. 1673 genes of all genes selected are unique to MAST. **[B]** and **[C]** The distribution of the expression fold-change and specificity scores for genes identified by the different methods. *getPValues()* selects a set of genes that enrich for high expression fold-change **[B]** and high specificity scores **[C]**. **[D]** The distribution of the posterior probability of observing the cluster given that the gene was observed (equation 1, main text) for genes identified by the different methods. MAST seems to weigh higher posterior probability positively in determining whether a gene is differentially expressed, even if fold-change ratio is relatively low, which perhaps explains its uniquely high sensitivity compared to genesorteR and Wilcoxon test. Data used here is the 10x mouse embryonic heart data (see Supplementary Text 2). "All Selected Genes" is the union of all genes selected by all methods.

Fig. S8

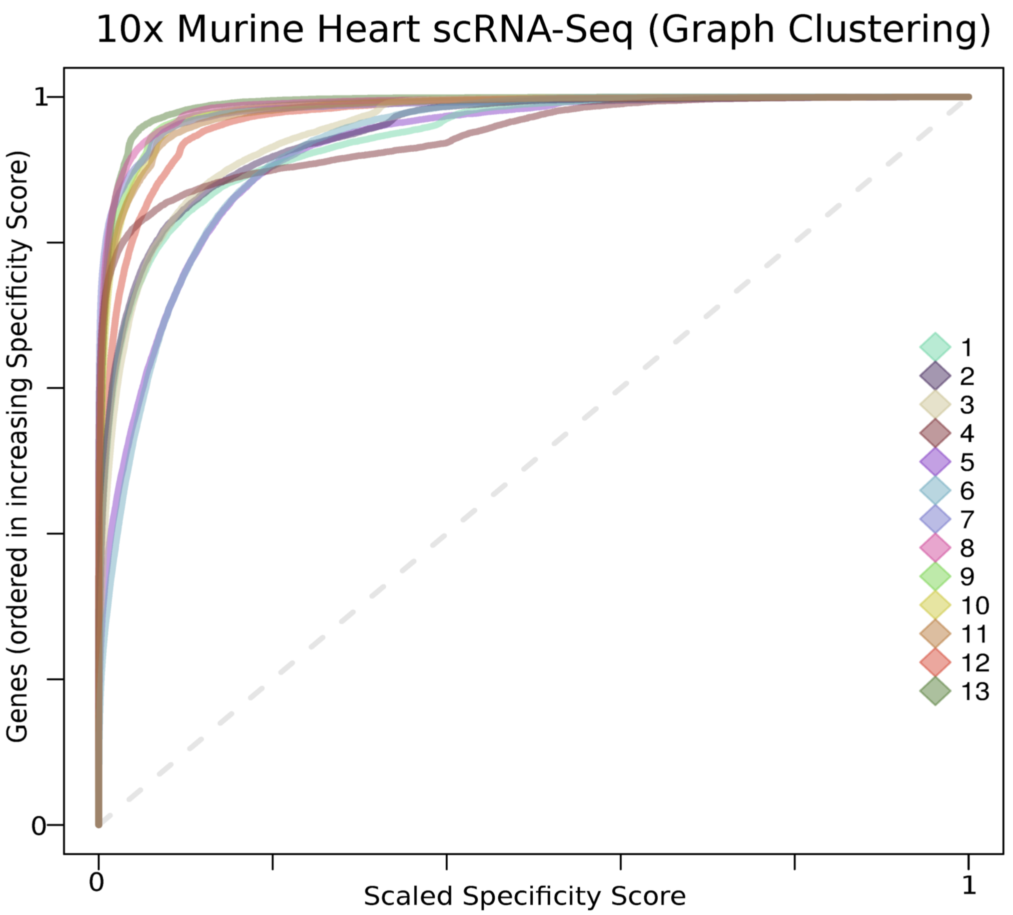
10x Murine Heart scRNA-Seq (Graph Clustering)
Genes (ordered in increasing Specificity Score)
Scaled Specificity Score
0
1
0
1
1
2
3
4
5
6
7
8
9
10
11
12
13

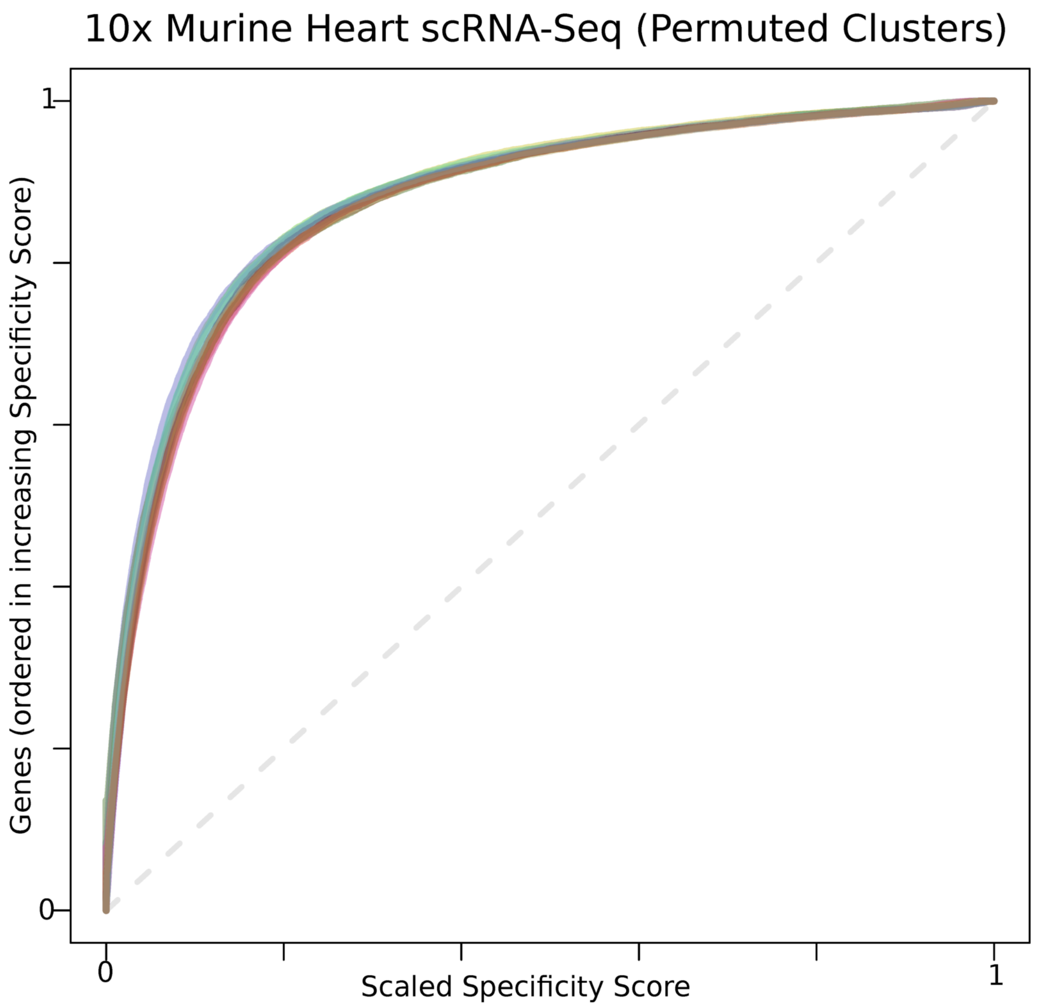
10x Murine Heart scRNA-Seq (Permuted Clusters)
Genes (ordered in increasing Specificity Score)
Scaled Specificity Score
0
1
0
1

**Figure S8.** *getClassAUC()* allows for assessing clustering quality by sorting genes by their specificity score in each cluster. Curves for well-separated clusters are further away from the diagonal, while curves for lower quality clusters are closer to the diagonal. The Area Under the Curve (AUC, main Figure 1b) can be used to quantify clustering quality. **Top:** the plot produced by getClassAUC on the 10x embryonic mouse heart data using graph clustering (see main Figure 1a, see Supplementary Text 2). **Bottom:** the same plot but using 13 randomly permuted clusters, signifying the "background" curves expected by random clustering of this dataset. Different datasets might have different random clustering curves. This metric is a relative metric related to the gene expression matrix at hand rather than an absolute clustering quality metric.

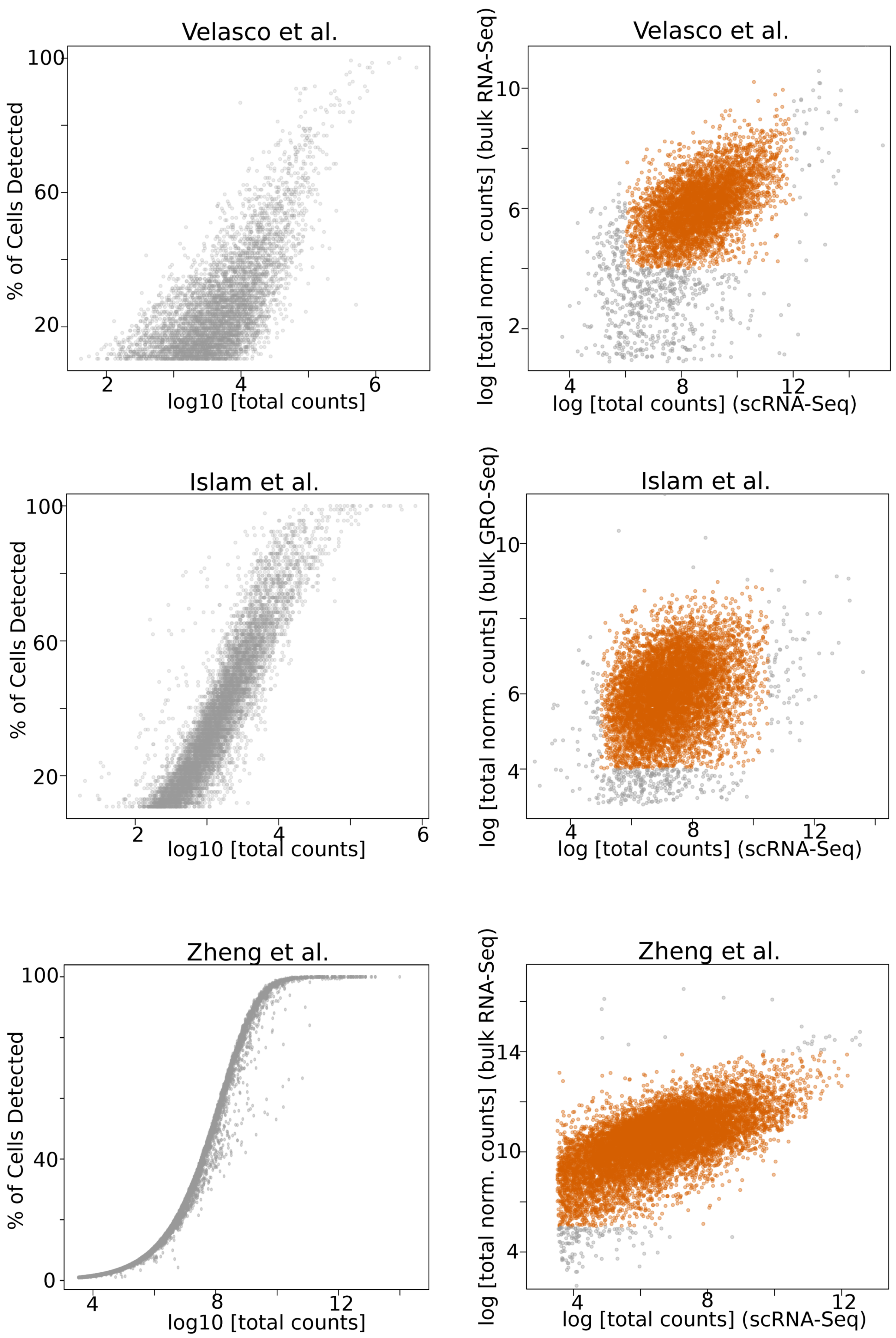
Velasco et al.
% of Cells Detected
log10 [total counts]
Velasco et al.
log [total norm. counts] (bulk RNA-Seq)
log [total counts] (scRNA-Seq)
Islam et al.
% of Cells Detected
log10 [total counts]
Islam et al.
log [total norm. counts] (bulk GRO-Seq)
log [total counts] (scRNA-Seq)
Zheng et al.
% of Cells Detected
log10 [total counts]
Zheng et al.
log [total norm. counts] (bulk RNA-Seq)
log [total counts] (scRNA-Seq)

Fig. S9

**Figure S9.** Characteristics of data used for benchmarking differential expression methods for scRNA-Seq. **Left:** Total counts versus detection rate (see also Supplementary Figures S1 and S2) for all genes show that the two variables are related in all three data sets but to varying extents. **Right:** Correspondance between the total expression for all genes in the scRNA-Seq data and the bulk expression data used for benchmarking. Only genes highlighted in orange were selected for benchmarking analyses in order to minimize benchmarking errors due to differences between the single cell and bulk data.

Fig. S10

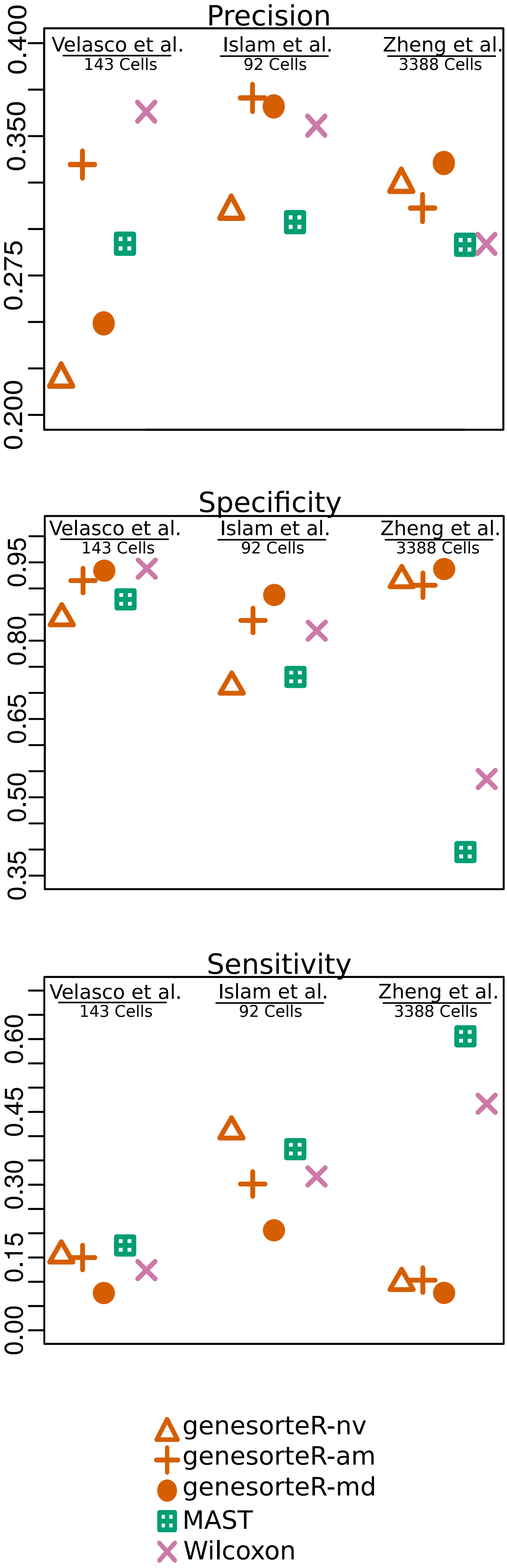

Precision
Velasco et al.
143 Cells
Islam et al.
92 Cells
Zheng et al.
3388 Cells
0.200
0.275
0.350
0.400
Specificity
0.35
0.50
0.65
0.80
0.95
Sensitivity
0.00
0.15
0.30
0.45
0.60
genesorteR-nv
genesorteR-am
genesorteR-md
MAST
Wilcoxon

**Figure S10.** Precision, specificity and sensitivity analysis for scRNA-Seq marker gene detection methods. nv=naive binarization, am=adaptive median binarization and md=median binarization. See also Main Figure 2.

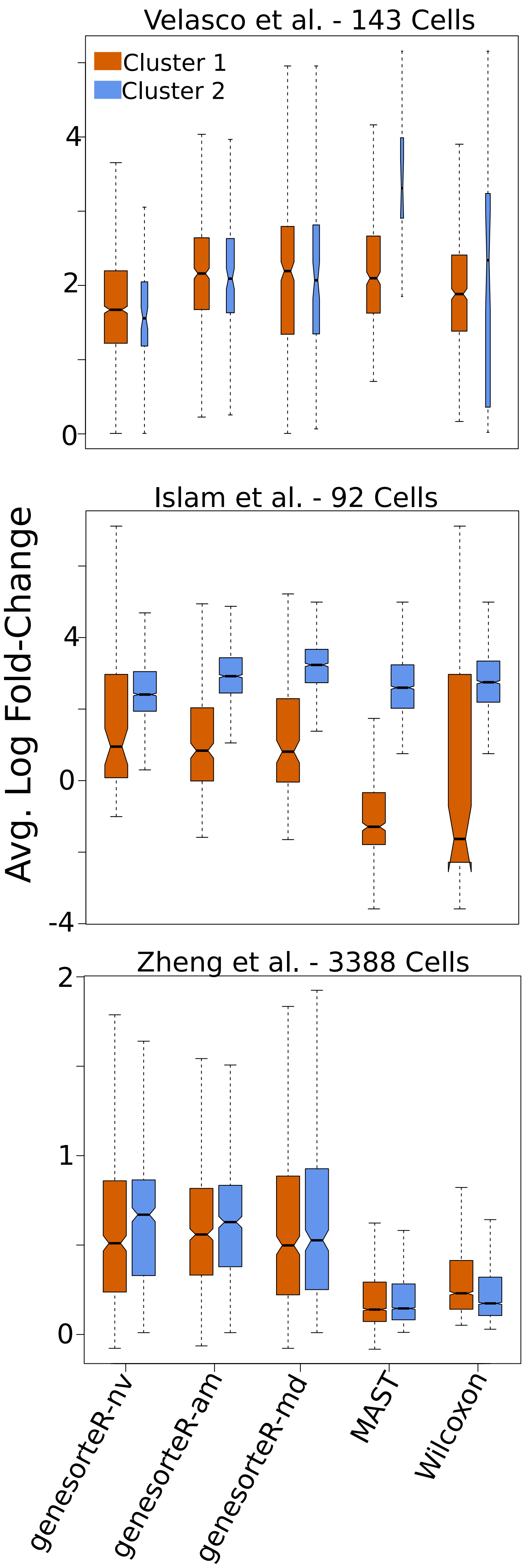
Velasco et al. - 143 Cells
Cluster 1
Cluster 2
Islam et al. - 92 Cells
Zheng et al. - 3388 Cells
Avg. Log Fold-Change
0
2
4
-4
1
genesorteR-nv
genesorteR-am
genesorteR-md
MAST
Wilcoxon

Fig. S11

**Figure S11.** Average log-fold change for genes detected by different scRNA-Seq marker detection methods, according to the “second formula” (mean[log[expression(pos. cluster)+1]] - mean[log[expression(neg. Cluster)+1]], see Results section in the Main text). Negative average log-fold change values are possible because genes were thresholded on positive log-fold change using the “first formula”. See also Main Figure 3 and Results section in the Main text. genesorteR’s *getTable()* function reports average log-fold change values using the “second formula”.

### Fig. S12

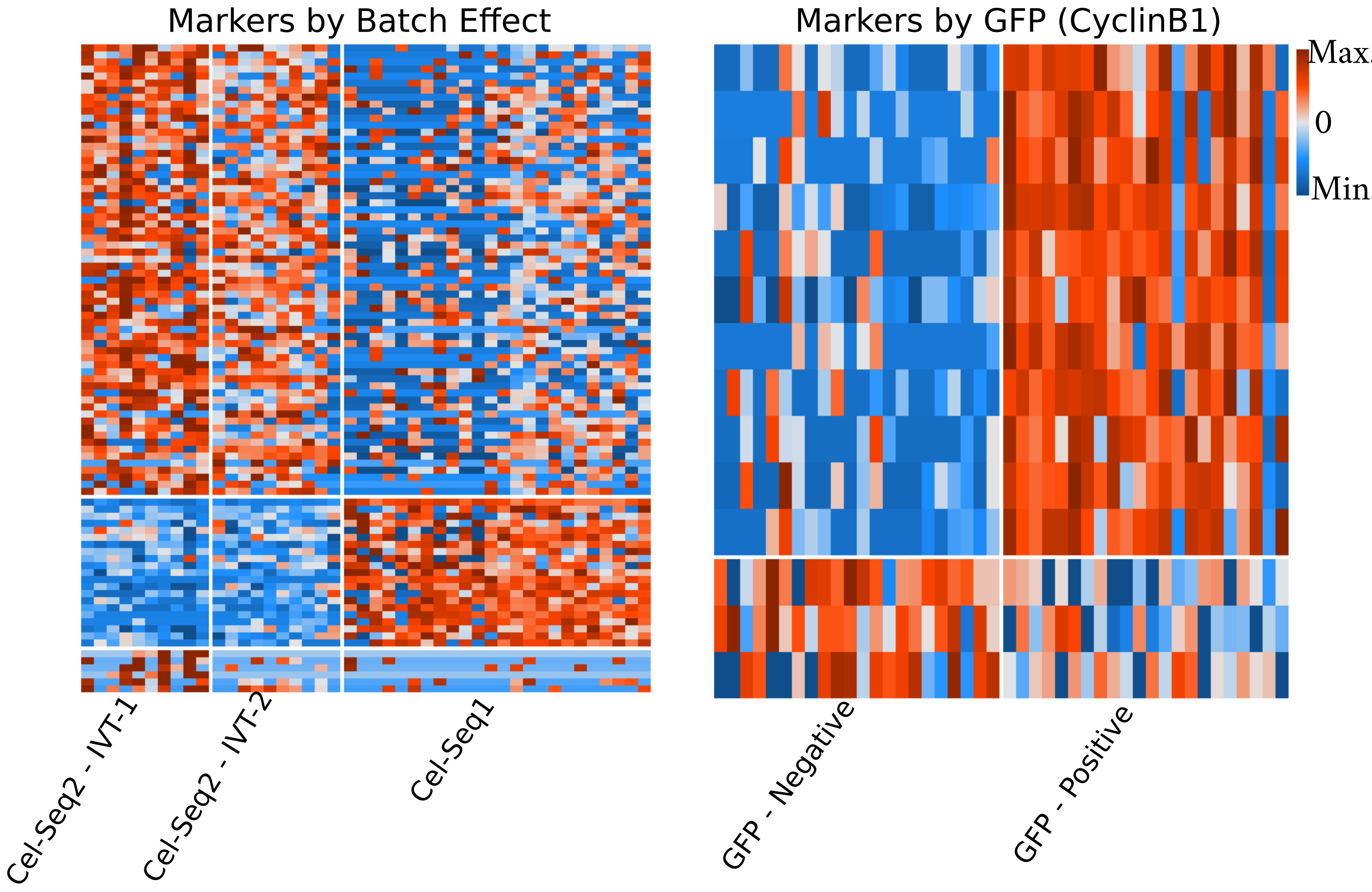

Markers by Batch Effect
Cel-Seq2 - IVT-1
Cel-Seq2 - IVT-2
Cel-Seq1
Markers by GFP (CyclinB1)
GFP - Negative
GFP - Positive
Max.
0
Min

**Figure S12.** Using genesorteR to find markers in CEL-Seq2 data (see Supplemetary Text 2). Markers that discriminate between technical batches (left) and between CyclinB1 positive and negative cells (right). Markers were identified using *getMarkers()* and plotted and clustered using *plotMarkerHeat()*.

Fig. S13

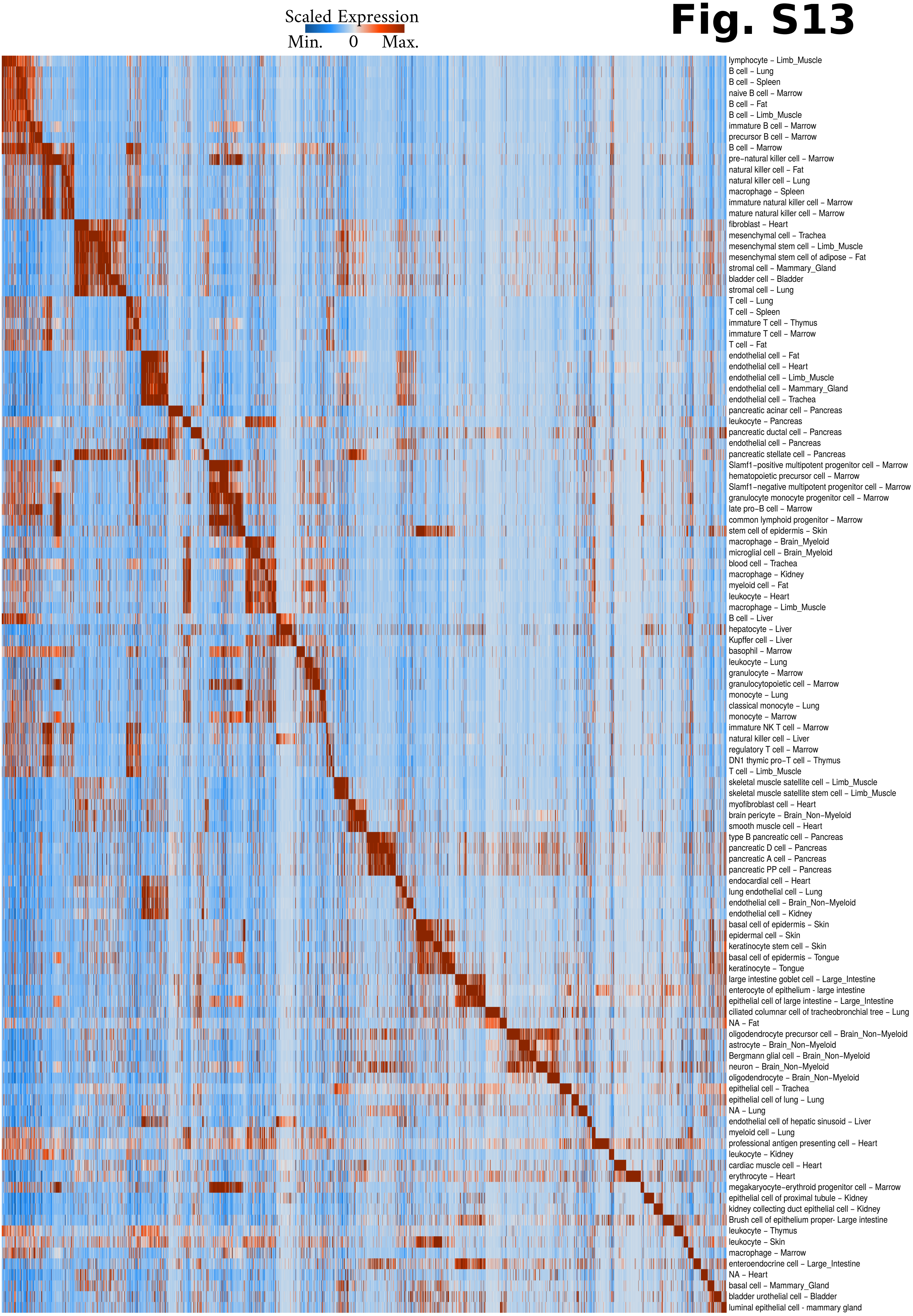
Scaled Expression
Min.
0
Max.
lymphocyte – Limb_Muscle
B cell – Lung
B cell – Spleen
naive B cell – Marrow
B cell – Fat
B cell – Limb_Muscle
immature B cell – Marrow
precursor B cell – Marrow
B cell – Marrow
pre−natural killer cell – Marrow
natural killer cell – Fat
natural killer cell – Lung
macrophage – Spleen
immature natural killer cell – Marrow
mature natural killer cell – Marrow
fibroblast – Heart
mesenchymal cell – Trachea
mesenchymal stem cell – Limb_Muscle
mesenchymal stem cell of adipose – Fat
stromal cell – Mammary_Gland
bladder cell – Bladder
stromal cell – Lung
T cell – Lung
T cell – Spleen
immature T cell – Thymus
immature T cell – Marrow
T cell – Fat
endothelial cell – Fat
endothelial cell – Heart
endothelial cell – Limb_Muscle
endothelial cell – Mammary_Gland
endothelial cell – Trachea
pancreatic acinar cell – Pancreas
leukocyte – Pancreas
pancreatic ductal cell – Pancreas
endothelial cell – Pancreas
pancreatic stellate cell – Pancreas
Slamf1−positive multipotent progenitor cell – Marrow
hematopoietic precursor cell – Marrow
Slamf1−negative multipotent progenitor cell – Marrow
granulocyte monocyte progenitor cell – Marrow
late pro−B cell – Marrow
common lymphoid progenitor – Marrow
stem cell of epidermis – Skin
macrophage – Brain_Myeloid
microglial cell – Brain_Myeloid
blood cell – Trachea
macrophage – Kidney
myeloid cell – Fat
leukocyte – Heart
macrophage – Limb_Muscle
B cell – Liver
hepatocyte – Liver
Kupffer cell – Liver
basophil – Marrow
leukocyte – Lung
granulocyte – Marrow
granulocytopoietic cell – Marrow
monocyte – Lung
classical monocyte – Lung
monocyte – Marrow
immature NK T cell – Marrow
natural killer cell – Liver
regulatory T cell – Marrow
DN1 thymic pro−T cell – Thymus
T cell – Limb_Muscle
skeletal muscle satellite cell – Limb_Muscle
skeletal muscle satellite stem cell – Limb_Muscle
myofibroblast cell – Heart
brain pericyte – Brain_Non−Myeloid
smooth muscle cell – Heart
type B pancreatic cell – Pancreas
pancreatic D cell – Pancreas
pancreatic A cell – Pancreas
pancreatic PP cell – Pancreas
endocardial cell – Heart
lung endothelial cell – Lung
endothelial cell – Brain_Non−Myeloid
endothelial cell – Kidney
basal cell of epidermis – Skin
epidermal cell – Skin
keratinocyte stem cell – Skin
basal cell of epidermis – Tongue
keratinocyte – Tongue
large intestine goblet cell – Large_Intestine
enterocyte of epithelium - large intestine
epithelial cell of large intestine – Large_Intestine
ciliated columnar cell of tracheobronchial tree – Lung
NA – Fat
oligodendrocyte precursor cell – Brain_Non−Myeloid
astrocyte – Brain_Non−Myeloid
Bergmann glial cell – Brain_Non−Myeloid
neuron – Brain_Non−Myeloid
oligodendrocyte – Brain_Non−Myeloid
epithelial cell – Trachea
epithelial cell of lung – Lung
NA – Lung
endothelial cell of hepatic sinusoid – Liver
myeloid cell – Lung
professional antigen presenting cell – Heart
leukocyte – Kidney
cardiac muscle cell – Heart
erythrocyte – Heart
megakaryocyte−erythroid progenitor cell – Marrow
epithelial cell of proximal tubule – Kidney
kidney collecting duct epithelial cell – Kidney
Brush cell of epithelium proper- Large intestine
leukocyte – Thymus
leukocyte – Skin
macrophage – Marrow
enteroendocrine cell – Large_Intestine
NA – Heart
basal cell – Mammary_Gland
bladder urothelial cell – Bladder
luminal epithelial cell - mammary gland

**Figure S13.** Using genesorteR to rank genes for each cell type identified in the Tabula Muris Smart-Seq2 data (see Supplementary Text 2). *plotTopMarkerHeat()* was used to process the output of *sortGenes()* in order to produce a heatmap of scaled gene expression for top marker genes in each cluster. All cells in each cluster are averaged so that each cluster occupies one row. Each column displays one gene. The top 100 genes in each cluster are displayed, but each gene is plotted only once and is not re-plotted if it had already appeared in a “previous” cluster. Generally, we observe that cells cluster by phenotype (endothelial, epithelial...etc.) rather than by tissue of origin.

### Fig. S14

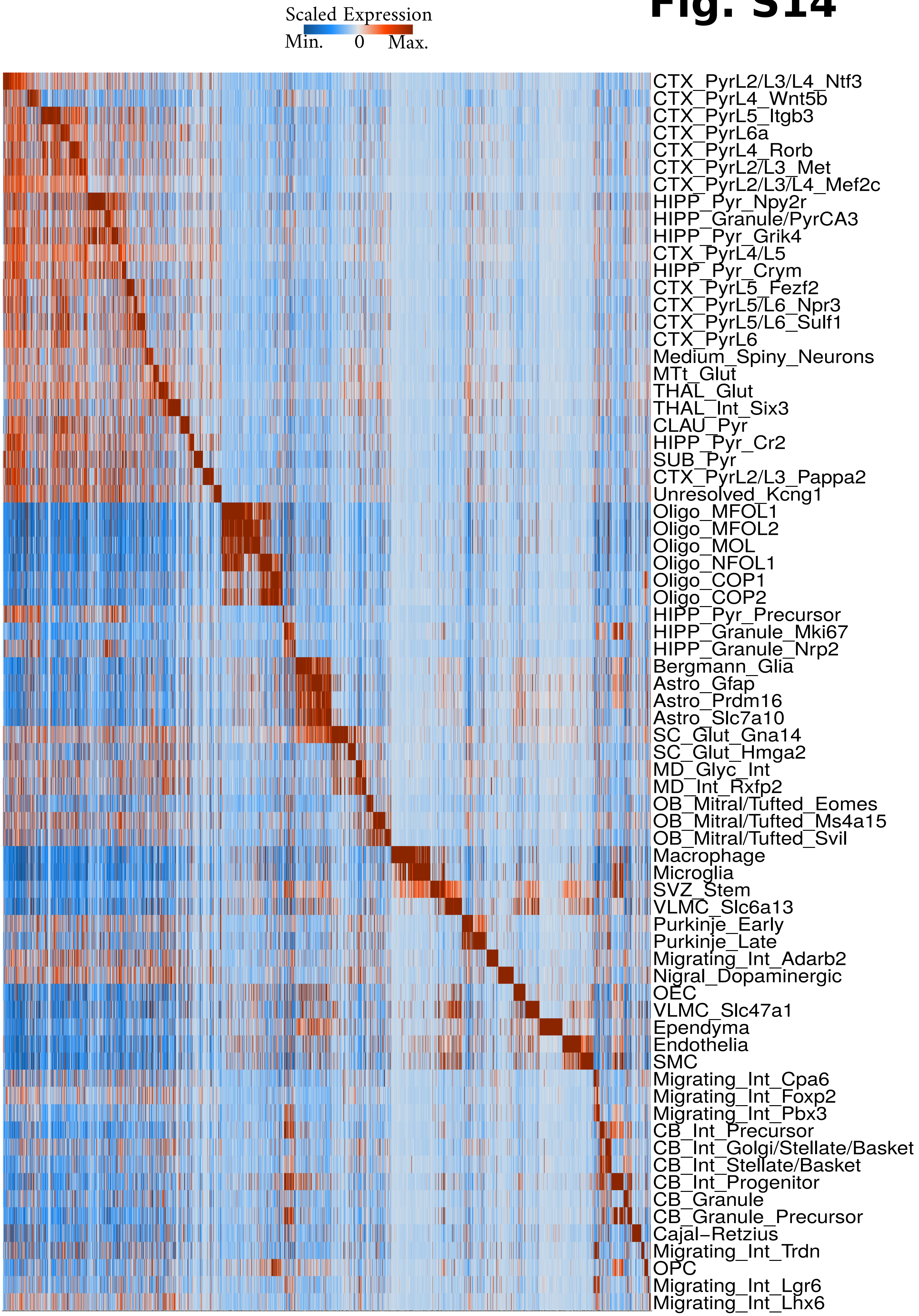
Scaled Expression
Min.
0
Max.
CTX_PyrL2/L3/L4_Ntf3
CTX_PyrL4_Wnt5b
CTX_PyrL5_Itgb3
CTX_PyrL6a
CTX_PyrL4_Rorb
CTX_PyrL2/L3_Met
CTX_PyrL2/L3/L4_Mef2c
HIPP_Pyr_Npy2r
HIPP_Granule/PyrCA3
HIPP_Pyr_Grik4
CTX_PyrL4/L5
HIPP_Pyr_Crym
CTX_PyrL5_Fezf2
CTX_PyrL5/L6_Npr3
CTX_PyrL5/L6_Sulf1
CTX_PyrL6
Medium_Spiny_Neurons
MTt_Glut
THAL_Glut
THAL_Int_Six3
CLAU_Pyr
HIPP_Pyr_Cr2
SUB_Pyr
CTX_PyrL2/L3_Pappa2
Unresolved_Kcng1
Oligo_MFOL1
Oligo_MFOL2
Oligo_MOL
Oligo_NFOL1
Oligo_COP1
Oligo_COP2
HIPP_Pyr_Precursor
HIPP_Granule_Mki67
HIPP_Granule_Nrp2
Bergmann_Glia
Astro_Gfap
Astro_Prdm16
Astro_Slc7a10
SC_Glut_Gna14
SC_Glut_Hmga2
MD_Glyc_Int
MD_Int_Rxfp2
OB_Mitral/Tufted_Eomes
OB_Mitral/Tufted_Ms4a15
OB_Mitral/Tufted_Svil
Macrophage
Microglia
SVZ_Stem
VLMC_Slc6a13
Purkinje_Early
Purkinje_Late
Migrating_Int_Adarb2
Nigral_Dopaminergic
OEC
VLMC_Slc47a1
Ependyma
Endothelia
SMC
Migrating_Int_Cpa6
Migrating_Int_Foxp2
Migrating_Int_Pbx3
CB_Int_Precursor
CB_Int_Golgi/Stellate/Basket
CB_Int_Stellate/Basket
CB_Int_Progenitor
CB_Granule
CB_Granule_Precursor
Cajal−Retzius
Migrating_Int_Trdn
OPC
Migrating_Int_Lgr6
Migrating_Int_Lhx6

**Figure S14.** The same plot as in Figure S13 but the mouse CNS SPLiT-Seq data (see Supplementary Text 2). This plot enables quick assessment of cell clustering: almost all cell clusters identified by *Rosenberg et al.* are relatively well-separated but clusters assigned to different layers of the cortex (labelled “CTX”, top) are somewhat similar to each other.

Fig. S15

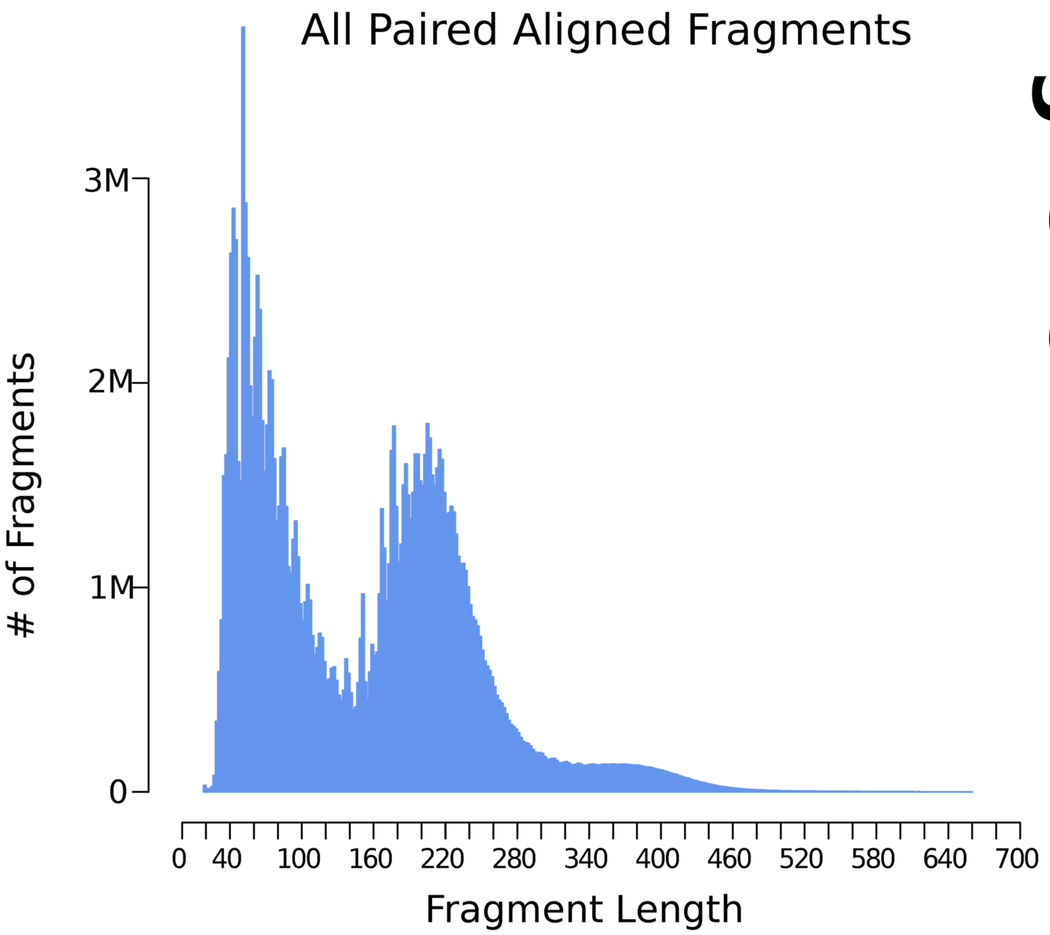
All Paired Aligned Fragments
### of Fragments
3M
2M
1M
0
0
40
100
160
220
280
340
400
460
520
580
640
700
Fragment Length

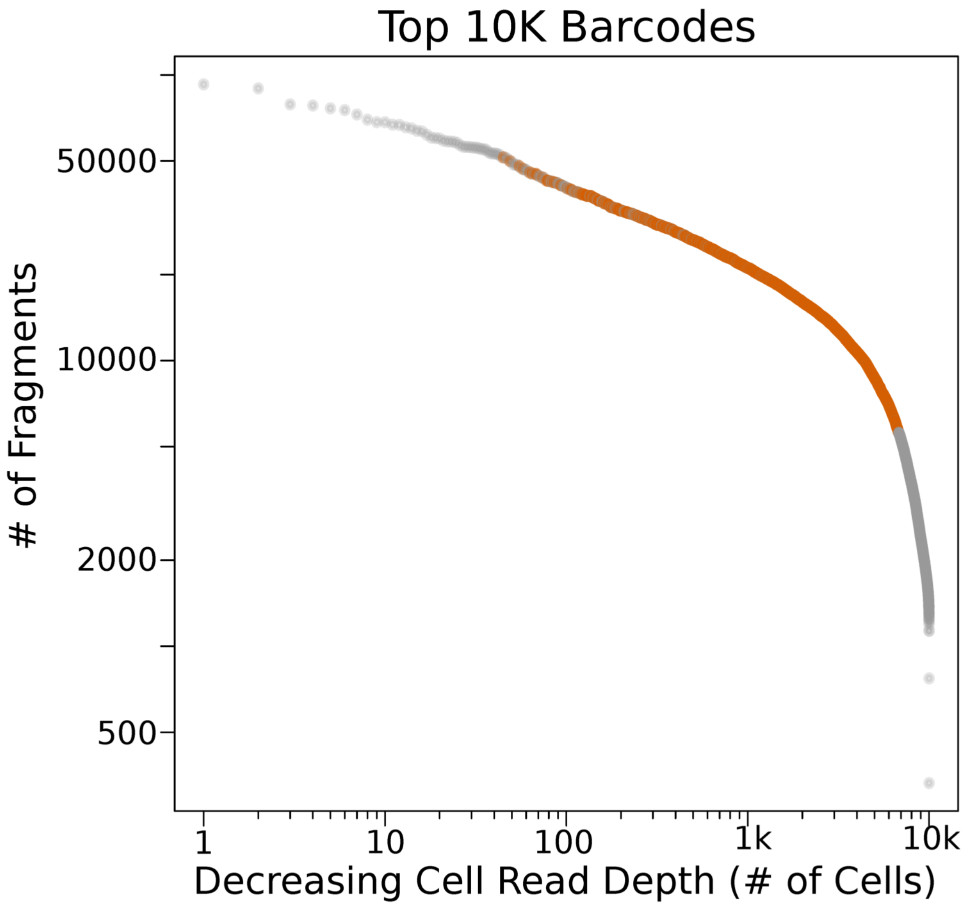
Top 10K Barcodes
### of Fragments
50000
10000
2000
500
1
10
100
1k
10k
Decreasing Cell Read Depth (# of Cells)

**Figure S15.** Characteristics of the scATAC-Seq data set reported in this manuscript. **Top:** Fragment length distribution of all aligned properly paired fragments excluding mitochondrial reads. **Bottom:** Cells arranged by total number of fragments assigned after alignment and filtering mitochnodrial reads. Only the top 10k cells by the number of fragments are shown. Cell highlighted in orange were selected for further analysis (see Supplementary Text 2).

### Fig. S16

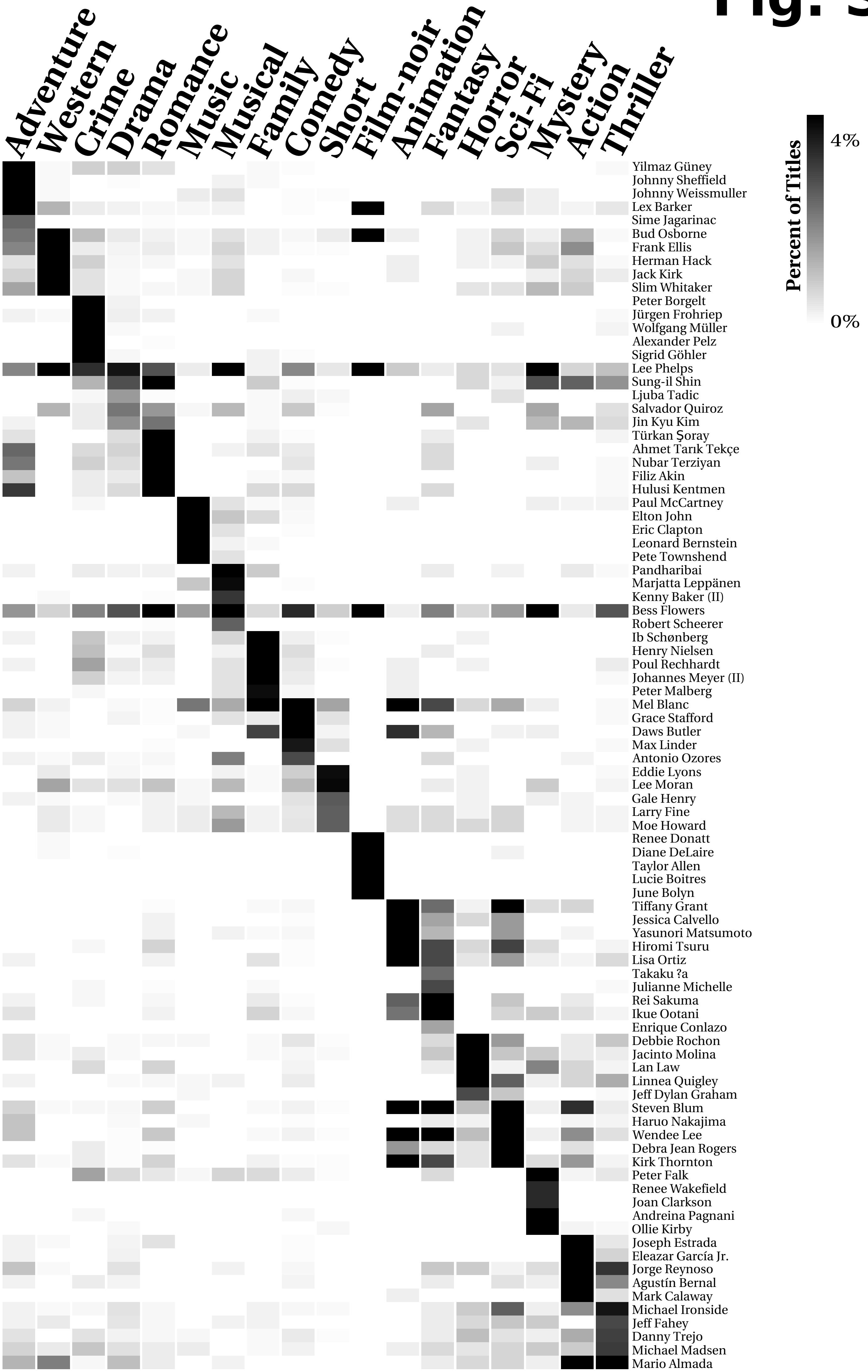

Adventure
Western
Crime
Drama
Romance
Music
Musical
Family
Comedy
Short
Film-noir
Animation
Fantasy
Horror
Sci-Fi
Mystery
Action
Thriller
Percent of Titles
4%
0%
Yilmaz Güney
Johnny Sheffield
Johnny Weissmuller
Lex Barker
Sime Jagarinac
Bud Osborne
Frank Ellis
Herman Hack
Jack Kirk
Slim Whitaker
Peter Borgelt
Jürgen Frohriep
Wolfgang Müller
Alexander Pelz
Sigrid Göhler
Lee Phelps
Sung-il Shin
Ljuba Tadic
Salvador Quiroz
Jin Kyu Kim
Türkan Şoray
Ahmet Tarık Tekçe
Nubar Terziyan
Filiz Akin
Hulusi Kentmen
Paul McCartney
Elton John
Eric Clapton
Leonard Bernstein
Pete Townshend
Pandharibai
Marjatta Leppänen
Kenny Baker (II)
Bess Flowers
Robert Scheerer
Ib Schønberg
Henry Nielsen
Poul Rechhardt
Johannes Meyer (II)
Peter Malberg
Mel Blanc
Grace Stafford
Daws Butler
Max Linder
Antonio Ozores
Eddie Lyons
Lee Moran
Gale Henry
Larry Fine
Moe Howard
Renee Donatt
Diane DeLaire
Taylor Allen
Lucie Boitres
June Bolyn
Tiffany Grant
Jessica Calvello
Yasunori Matsumoto
Hiromi Tsuru
Lisa Ortiz
Takaku ?a
Julianne Michelle
Rei Sakuma
Ikue Ootani
Enrique Conlazo
Debbie Rochon
Jacinto Molina
Lan Law
Linnea Quigley
Jeff Dylan Graham
Steven Blum
Haruo Nakajima
Wendee Lee
Debra Jean Rogers
Kirk Thornton
Peter Falk
Renee Wakefield
Joan Clarkson
Andreina Pagnani
Ollie Kirby
Joseph Estrada
Eleazar García Jr.
Jorge Reynoso
Agustín Bernal
Mark Calaway
Michael Ironside
Jeff Fahey
Danny Trejo
Michael Madsen
Mario Almada

**Figure S16.** Analysis of historical data from the Internet Movie Database (www.imdb.com, see Supplementary Text 2). The function *plotTopBinaryHeat()* was used to process the output of *sortGenes()* and produce a plot of the percentage of titles in which an actor appeared in each genre. All titles in each genre were averaged per actor such that each genre occupies only one column. The top 5 actors for each genre are shown.

### Fig. S17

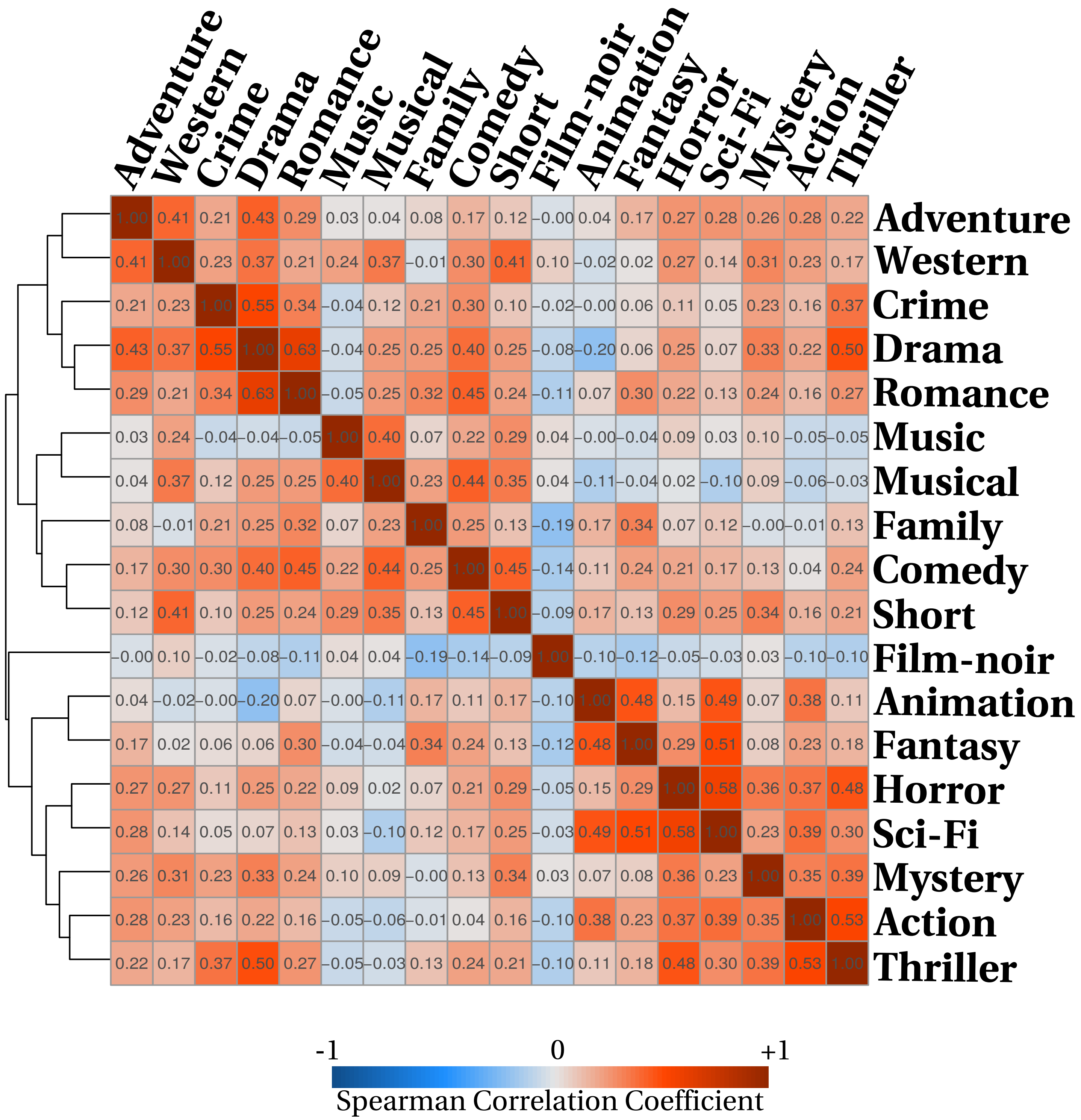
Adventure
Western
Crime
Drama
Romance
Music
Musical
Family
Comedy
Short
Film-noir
Animation
Fantasy
Horror
Sci-Fi
Mystery
Action
Thriller
1.00 0.41 0.21 0.43 0.29 0.03 0.04 0.08 0.17 0.12 −0.00 0.04 0.17 0.27 0.28 0.26 0.28 0.22 Adventure
0.41 1.00 0.23 0.37 0.21 0.24 0.37 −0.01 0.30 0.41 0.10 −0.02 0.02 0.27 0.14 0.31 0.23 0.17 Western
0.21 0.23 1.00 0.55 0.34 −0.04 0.12 0.21 0.30 0.10 −0.02 −0.00 0.06 0.11 0.05 0.23 0.16 0.37 Crime
0.43 0.37 0.55 1.00 0.63 −0.04 0.25 0.25 0.40 0.25 −0.08 −0.20 0.06 0.25 0.07 0.33 0.22 0.50 Drama
0.29 0.21 0.34 0.63 1.00 −0.05 0.25 0.32 0.45 0.24 −0.11 0.07 0.30 0.22 0.13 0.24 0.16 0.27 Romance
0.03 0.24 −0.04 −0.04 −0.05 1.00 0.40 0.07 0.22 0.29 0.04 −0.00 −0.04 0.09 0.03 0.10 −0.05 −0.05 Music
0.04 0.37 0.12 0.25 0.25 0.40 1.00 0.23 0.44 0.35 0.04 −0.11 −0.04 0.02 −0.10 0.09 −0.06 −0.03 Musical
0.08 −0.01 0.21 0.25 0.32 0.07 0.23 1.00 0.25 0.13 −0.19 0.17 0.34 0.07 0.12 −0.00 −0.01 0.13 Family
0.17 0.30 0.30 0.40 0.45 0.22 0.44 0.25 1.00 0.45 −0.14 0.11 0.24 0.21 0.17 0.13 0.04 0.24 Comedy
0.12 0.41 0.10 0.25 0.24 0.29 0.35 0.13 0.45 1.00 −0.09 0.17 0.13 0.29 0.25 0.34 0.16 0.21 Short
−0.00 0.10 −0.02 −0.08 −0.11 0.04 0.04 −0.19 −0.14 −0.09 1.00 −0.10 −0.12 −0.05 −0.03 0.03 −0.10 −0.10 Film-noir
0.04 −0.02 −0.00 −0.20 0.07 −0.00 −0.11 0.17 0.11 0.17 −0.10 1.00 0.48 0.15 0.49 0.07 0.38 0.11 Animation
0.17 0.02 0.06 0.06 0.30 −0.04 −0.04 0.34 0.24 0.13 −0.12 0.48 1.00 0.29 0.51 0.08 0.23 0.18 Fantasy
0.27 0.27 0.11 0.25 0.22 0.09 0.02 0.07 0.21 0.29 −0.05 0.15 0.29 1.00 0.58 0.36 0.37 0.48 Horror
0.28 0.14 0.05 0.07 0.13 0.03 −0.10 0.12 0.17 0.25 −0.03 0.49 0.51 0.58 1.00 0.23 0.39 0.30 Sci-Fi
0.26 0.31 0.23 0.33 0.24 0.10 0.09 −0.00 0.13 0.34 0.03 0.07 0.08 0.36 0.23 1.00 0.35 0.39 Mystery
0.28 0.23 0.16 0.22 0.16 −0.05 −0.06 −0.01 0.04 0.16 −0.10 0.38 0.23 0.37 0.39 0.35 1.00 0.53 Action
0.22 0.17 0.37 0.50 0.27 −0.05 −0.03 0.13 0.24 0.21 −0.10 0.11 0.18 0.48 0.30 0.39 0.53 1.00 Thriller
-1
0
+1
Spearman Correlation Coefficient

**Figure S17.** Heirarchical clustering of genres based on the correlation coefficients between actor specificity scores (the output of *sortGenes()*) for each genre. Specificity score for the top 20 actors of each genre were used. The function *plotCorrelationHeat()* was used to produce this plot.

#### genesorteR: Feature Ranking in Clustered Single Cell Data

### Supplementary Text 1

In the section, we summarize the current functionality of genesorteR and expand on the discussion of some of its methods. Note that symbol definition here follows the same convention as in the main text.

- **sortGenes()**

*sortGenes* is the main function of the genesorteR package. It takes a gene expression matrix and cell cluster assignments. It binarizes the expression matrix and calculates empirical statistics on gene expression in each cluster.

When *binarizeMethod* is "median", expression matrix binarization is done by estimating a cut-off value uniformly on all values in the matrix, which is equal to the median of all non-zero entries in the matrix. When *binarizeMethod* is "naive", all non-zero entries are kept. Enforcing further sparsity on the matrix using the “median” method is recommended in most cases as it enables better separation between clusters by removing “spurious” detection of genes at a very low expression level. However, we find that keeping the matrix as is also works well and the decision to enforce further sparsity or not may be a dataset-specific decision.

The specificity score matrix is returned by *sortGenes* as “specScore”. It balances the posterior probability of observing a cell cluster given the gene (a measure of gene specificity) with the gene conditional probability given the cluster (a measure of gene expression level, see Equations 1 and 2, Main Text). This ensures that highly specific genes are also highly expressed. The “specScore” matrix is considered the main output of this function, and on which many of the remaining calculations by other functions in the genesorteR package are performed. The values in this matrix can be used to rank features (genes in scRNA-Seq) in cell clusters.

*sortGenes* can in principle be applied to both a raw count matrix or a normalized log-count expression matrix.

- **getMarkers()**

*getMarkers* processes the output of *sortGenes* to select relatively small sets of marker genes. The idea is to enable cell type validation experiments by proposing small sets of markers that might be used to identify cell types uniquely in wet-lab experiments. *getMarkers* relies on calculating an entropy-like metric (called here Gene Specificity Shannon Index, see Main Text) calculated on the scaled gene-cluster specificity score. The intuition is that genes with low Gene Specificity Shannon Index are either lowly expressed or highly specific to one or few cell clusters.

Therefore, we select the top $n\%$ of genes according to the scaled specificity score and then cluster those genes based on their Gene Specificity Shannon Index. The scaled specificity score for gene $g_j$ in cluster $t_i$ is calculated as:

$$sc_{ij} = {s_{ij} - \min(\overrightarrow{S_{t_i}})} \Big/ {\max(\overrightarrow{S_{t_i}}) - \min(\overrightarrow{S_{t_i}})}$$

Where $s_{ij}$ is the specificity score for gene $g_j$ in cluster $t_i$ and $\overrightarrow{S_{t_i}}$ is the vector of specificity scores for all $m$ genes in cluster $t_i$. Scaling the specificity score is done to try to guarantee that marker genes will be selected for each cluster even if some of the clusters are not well separated.

The percent of genes to select for clustering ($n$) is controlled by the user and is 1% by default.

*getMarkers* can also return the mutual information between gene expression for gene $j$ ($m_j$) and cell clusters:

$$m_j = H(\overrightarrow{g_j}) + H(\vec{t}) - H(\overrightarrow{g_j}, \vec{t})$$

Where $H(\overrightarrow{g_j})$ is the entropy of the binarized expression vector for gene $j$, $H(\vec{t})$ is the entropy of the cell cluster identity vector and $H(\overrightarrow{g_j}, \vec{t})$ is the joint entropy of $\overrightarrow{g_j}$ and $\vec{t}$. Therefore, $m_j$ can be seen as a specificity metric to rank genes overall by the variability of their gene expression among cell clusters.

*getMarkers* can also return a Cluster Specificity Shannon Index for each cell cluster $t_i$, indicating how well identified this cell cluster is:

$$SI_i = -\sum_{i=1}^{i=l} sc_{ij} \,.\log(sc_{ij}) \qquad i \in \vec{M}$$

Where $\vec{M}$ is a vector of length $l$ containing indeces of genes selected by *getMarkers* as marker genes. Cluster Specificity Shannon Index is calculated on the scaled specificity score considering only marker genes selected by *getMarkers.* Therefore, it answers the question: how well-separated this cell cluster is given the marker genes selected by *getMarkers.* It is often interesting to compare this with the more global cluster quality metric returned by *getClassAUC* (see below).

- **<u>plotMarkerScores()</u>**

*plotMarkerScores* plots scatter plots of the specificity score and the gene specificity Shannon index to investigate *getMarkers* results.

- **<u>getPValues()</u>**

*getPValues* performs a permutation test on the gene-cell type specificity score to obtain a *p*-value on each gene-cell type specificity score. The permutation keeps everything the same except that cell type assignments are permuted between the cells (but note that cell type proportions are also kept the same). This function can be a way to select all differentially expressed genes between all cell classes globally in a dataset in one go or to define all statistically significantly differentially expressed genes for each cluster. In a way, it also asks given the gene expression data, how valid the clustering is. If clustering is poor, very few or no genes will be detected using this method.

The null distribution of specificity scores is calculated based on the specificity score matrices resulting from all performed permutations, but excluding zero specificity score. Excluding zeros is done because given a binarized gene expression matrix and a corresponding distribution of cell type identities, a certain proportion of zeros in the specificity score matrix is expected regardless of actual cell-cluster identity assignment. For most genes, the expected proportion of zeros can only be modelled and not exactly empirically calculated. Therefore, we found it impossible without further model-based assumptions to consider the expected "true" proportion of zeros in the specificity score matrices resulting from cell cluster permutations. Thus far, we decided to ignore all zeros when considering the null distribution of specificity scores arising from cell cluster permutations.

*p*-values for each gene-cluster are calculated from the specificity score null distribution based on the formula by *Phipson and Smyth*[1] for permutations without replacement. Note that not many permutations are required since the number of permuted values is typically relatively large: equal to $(m \,.\, k \,.\, p) - z$ , where $p$ is the number of permutations and $z$ is the number of zero entries in all permuted specificity score matrices. As noted by *Phipson and Smyth*[1], increasing the number of permutations increases the power to detect more genes at a given $\alpha$ level. The minimum possible *p*-value is equal to $1/[(m \,.\, k \,.\, p) - z + 1]$ .

- **getTable()**

gettable takes the output of *sortGenes* and *getPValues* and summarizes differential expression results for all clusters in one table. It additionally calculates average log-fold change and allows for gene thresholding based on log-fold change and adjusted p-values.

- **plotCorrelationHeat()**

*plotCorrelationHeat* uses the specificity scores from *sortGenes* to correlate cell clusters with each other and plots a heatmap of the correlations.

- **getClassAUC()**

getClassAUC implements a method to investigate clustering quality. It processes the output of *sortGenes* to obtain a curve for each cell cluster for all gene specificity scores against their ranking in the cluster. The Area Under the Curve (AUC) can be used as a measure of clustering quality in terms of the possibility to identify cell clusters.

The idea is that given the specificity score for all genes in a certain cell cluster, we can assume that a well-separated easily-identified cell cluster will have a relatively small number of genes that have a very high specificity score. Top marker genes for a cluster that is poorly separated from other cell clusters will have average or low specificity scores that are not highly different from the specificity scores of other genes for that cluster. Sorting the genes for each cell cluster by their specificity score and plotting the scaled scores in order creates a curve that should be far from the diagonal for well-separated clusters but close to the diagonal for poorly-separated clusters. The AUC of this curve can be used to quantify this intuition and estimate a clustering quality metric.

This method is inspired by ChIP-Seq quality metrics proposed in *Diaz et al.* [2] where the concentration of sequencing reads in relatively few genomic bins is an indicator of high-quality data.

- **plotMarkerHeat()**

*plotMarkerHeat* plots a heatmap of expression values for a select set of genes supplied by the user (for example the output of *getMarkers*), and optionally also clusters those genes. Clustering is done using k-means but on the Euclidean norm-scaled gene expression vectors (not binarized). If $E_j$ is the gene expression vector for gene $j$, then scaled is:

$$sE_j = {E_j}/{\|E_j\|_2}$$

This is inspired by the "cosine-normalization" proposed in *Haghverdi et al.* to normalize cell expression vectors[3]. We find this to be a quick and straightforward method to obtain stable gene clusters without the need for complicated clustering algorithms or data imputation.

By default, *plotMarkerHeat* plots every single cell in one column, this might not be practical for large data. Therefore, *plotMarkerHeat* also enables cell averaging to make plotting data with a large number of cells possible. For example, if "averageCells" is set to 10 every 10 cells will be averaged (without averaging across cell clusters) before plotting the heatmap.

- **plotTopMarkerHeat()**

*plotTopMarkerHeat* plots a heatmap of expression values for the top genes in each cluster, as defined by feature ranking via the specificity scores from *sortGenes*.

*plotTopMarkerHeat* is essentially a convenience wrapper around *plotMarkerHeat* that plots a heatmap of the top *n* genes (10 by default) in each cell cluster. Unlike *plotMarkerHeat, plotTopMarkerHeat* takes the output of *sortGenes* as the only required input. *plotTopMarkerHeat* cannot cluster genes, but can perform cell averaging.

Both *plotMarkerHeat* and *plotTopMarkerHeat* use the pheatmap R package for plotting heatmaps (https://CRAN.R-project.org/package=pheatmap).

- **plotBinaryHeat() and plotTopBinaryHeat()**

*plotBinaryHeat* and *plotTopBinaryHeat* are similar to *plotMarkerHeat* and *plotTopMarkerHeat* respectively but operate on binary expression matrices. When cell averaging is requested, the average is the fraction of cells where the features were detected.

**genesorteR: Feature Ranking in Clustered Single Cell Data**

### Supplementary Text 2

- **10X 10k Cells from Embryonic Mouse Whole Heart**

We started with the filtered gene/cell count matrix output provided by 10X Genomics (downloaded on April 30 2019)[1]. The matrix contained 7713 candidate cells. After ribosomal RNA genes and mitochondrial genes were removed, cells that had less than 1000 UMI counts were filtered out. Genes that were captured in less than 1% of all cells were filtered out. Finally, cells were filtered based on mitochondrial RNA content and bias toward highly expressed genes as follows: cells were clustered into two clusters using a bivariate Gaussian mixture with two components learned on log10(total UMI counts per cell excluding mitochondrial UMIs) and percent of mitochondrial UMI per cell. Clustering was performed using the R package Mclust v5.4.3 (parameters: modelNames="EII")[2]. Cells falling into the cluster with higher mitochondrial content cells were excluded. Simultaneously, cells whose total counts in the top 500 genes represented more than one and half times the median of total counts in the top 500 genes of all cells were removed (calculated cut-off was 94.7%). Finally, cells with more than 100,000 total UMI counts were removed. This filtering process resulted in 5070 cells, with a total of 13846 genes across all cells and an average of 3372 genes per cell.

Size factors for count normalization were determined using the Scran R package[3] function computeSumFactors v1.10.2 (parameters: sizes = seq(20, 200, 20), positive = FALSE)[4]. Variable genes were determined using the Scran package with an FDR cut-off of 0.001 (3382 genes). Cell expression vectors restricted to the variable genes were centred at the mean and Singular Value Decomposition was performed. The left singular vectors are "meta-genes" expressing a lower dimensional representation of the gene expression program in each cell[4]. We selected the top 16 meta-genes based on the knee of the curve of the singular values, which accounted for approximately 57.5% of the total variance present in the data. We used the top 16 meta-genes to generate the UMAP[5] cell projection using the R package umap v0.2 (parameters: min_dist=2, set_op_mix_ratio=1, local_connectivity=0.5, bandwidth=1, alpha=5, spread=10, https://github.com/tkonopka/umap).

To cluster cells into distinct cell types, we calculated the approximate k-nearest neighbours for each cell using the 16 top meta-genes based on their Euclidean distance using the function nn2 in the R package RANN v2.6.1 (parameters: searchtype = "priority", treetype = "bd", https://github.com/jefferis/RANN). The number of nearest neighbours "k" was set to the square root of the number of cells. We then sorted the distances between all k-nearest neighbours and kept the smallest *n* distances where n=[sqrt(number of cells) x number of cells]. Finally, we determined cell clusters using the Louvain graph clustering algorithm[6], as implemented in the igraph R package v1.2.4[7], on the undirected graph resulting from all the remaining *n* connections. We obtained 13 clusters from the Louvain algorithm (Figure 1a).

We ran sortGenes from the genesorteR package with default parameters on the normalized logcounts expression matrix and the cell clustering information. getMarkers was run with quant = 0.95, getPValues was run with numPerm = 20. For heatmaps using plotMarkerHeat and plotTopMarkerHeat, we averaged every 10 cells. Intersection between gene sets identified by Scran and genesorteR was visualized using the UpSetR R package[8].

- **CEL-Seq2 Data**

We started from gene expression count data for mouse with accession numbers GSE78779_CS1_manual and GSE78779_CS2_manual (24 cells using CEL-Seq1 protocol and 20 cells using CEL-Seq2 protocol)[9]. We analyzed ERCC transcripts and endogenous transcripts separately. For ERCC transcripts, we kept any gene that was detected in at least one cell. For endogenous transcripts, we kept any gene that was detected in at least 10% of the cells (11420 genes). No cell filtering was performed which resulted in 11420 genes detected across 44 cells, with an average of 7036 genes per cell.

The dataset contains two levels of clustering, one based on effects due to the difference between CEL-Seq1 and CEL-Seq2 including two CEL-Seq2 batches, and one based on CyclinB1 expression. We used genesorteR to detect differences along technical batches and along CyclinB1 expression. We used getMarkers setting quant to 0.95 to define potential marker genes across batch effect and along CyclinB1 expression.

- **Tabula Muris Smart-Seq2 Data**

Tabula Muris Smart-Seq2 data[10] was obtained using the TabulaMurisData R package (snapshot date: 2018-10-30)[11]. We analyzed ERCC and endogenous transcripts separately. Considering only endogenous genes, we kept cells that had at least 500 genes and 50000 reads, and whose top 500 genes accounted for less than 99% of all reads. Finally, we removed genes that were detected in less than 1% of all cells. This left 23341 genes across 43910 cells distributed over 115 clusters with an average of 3149 genes per cell. We normalized cells by the total number of reads and we used the resulting matrix to rank genes for each cell cluster using sortGenes with default parameters.

- **SPLiT-Seq Mouse CNS Data**

We started with the file GSM3017261_150000_CNS_nuclei.mat (GEO accession number: GSE110823)[12] and removed nuclei whose type was "Unresolved", leaving expression count data for 26894 genes across 95005 nuclei, with an average of 622 expressed genes per nucleus. Cell count data was normalized using total cell count size factors. Since this data set is relatively very sparse, sortGenes was run with binarizeMethod set to "naive"

- **10X Jurkat-HEK293T Data**

Data were processed and clustered in the same way as 10X mouse embryonic heart data (see above), with the exception that we used k-means to cluster cells based on the left singular vectors into 2 clusters. The filtered expression matrix was downloaded from 10x Genomics website on August 10 2019[13,14]. This resulted in 12828 genes across 3388 cells with an average of 3391 genes per cell.

To produce a ground truth dataset, public bulk RNA-Seq tag count data for Jurkat and HEK293T cell lines were download from the ARCHS4 database[15]. Samples that had a total count of less than 10^7 reads across all genes were removed, and genes that had less than 5 reads on average per sample were removed. Mitochondrial and ribosomal RNA and protein genes were removed. Data were normalized using the TMM method and differential expression was performed in the edgeR R package version 3.24.3[16]. p-values were adjusted using the Bonferroni method. Genes with an adjusted p-value < 0.01 and an absolute average log-fold change > 1 were considered as differentially expressed.

#### - EB and iSMN Data

Single cell gene expression counts were obtained from accession number GSE81275 (GSE81275_esc1.expMatrix.txt)[17]. Only 0h (Embryonic Bodies) and 48h (induced Spinal Motor Neurons) cells were considered. Mitochondrial genes and ribosomal genes were removed. Cells where less than 1% of genes were detected were removed and genes that were expressed in less than 10% of all cells were removed. Gene expression counts were normalized using the total cell count size factors.

To produce a ground truth dataset, we used bulk RNA-Seq data from the same conditions from the same publication[17]. Expression read counts were quantified from fastq files using the Gencode mouse transcriptome[18] version M13, and using RSEM[19] version 1.3.0 with default parameters and bowtie2[20] version 2.2.6 as an aligner. Genes that had less than 10 reads per sample on average were removed. Counts TMM normalization and differential expression were performed using the edgeR R package version 3.24.3[16]. p-values were adjusted using the Bonferroni method and genes with an adjusted p-value < 0.01 were considered as differentially expressed.

Only genes that were detected in both the single cell data and the bulk RNA-Seq data were kept resulting in 5548 genes across 143 cells with an average of 1476 genes per cell.

#### - mESC and MEF Data

Single cell gene expression counts were obtained from accession number GSE29087 (GSE29087_L139_expression_tab.txt)[21]. Only embryonic stem cells and mouse embryonic fibroblasts were considered. Mitochondrial and ribosomal genes were removed and genes that are detected in less than 10% of all cells were removed. Gene expression counts were normalized using total cell count size factors. This resulted in 7897 genes across 92 cells with average of 3052 genes per cell.

To obtain a ground truth dataset, we used bulk GRO-Seq data from the same two cell types[22]. Fastq files with SRA archive accession numbers SRR097863, SRR097864, SRR097862, SRR097858, SRR097859, SRR097860, SRR097861, SRR097854, SRR097855, SRR097856 and SRR097857 were aligned to the mm10 genome using bowtie2 version 2.2.6[20] and reads overlapping genes obtained from Gencode mouse transcriptome[18] annotation version M20 were counted in a stranded manner in both introns and exons to obtain gene expression counts. Genes that had less than 5 reads per sample on average were removed and ribosomal and mitochondrial genes were removed. Read count normalization using the TMM method and differential expression analysis were performed in the edgeR R package version 3.24.3[16]. Genes with a Bonferroni corrected p-value < 0.01 were considered as differentially expressed.

Only genes that were detected in both the single cell data and the bulk GRO-Seq data were kept resulting in 7897 genes across 92 cells with average of 3052 genes per cell for the single cell data.

#### - Comparison with MAST and the Wilcoxon Rank-sum Test

We used the Seurat R package (v3.0.1)[23] to run MAST[24] and perform Wilcoxon Rank-sum test[25] .The findAllMarkers function from Seurat was used setting pos.only to TRUE in order to obtain only positive markers, as this compares more closely to the behaviour of genesorteR's getPValues (see Supplementary Text 1). We determined the intersection between the genes detected by genesorteR getPValues (sortGenes was run with default parameters, getPValues was run with numPerm set to 20) and by Seurat using the UpSetR R package[8].

The maximum log fold change value for each gene (Supplementary Figure S7b) is the maximum value obtained from calculating the fold change of the average gene expression of cells in each cluster relative to all cells in all the other cluster. The maximum specificity score and maximum cluster posterior probability (Supplementary Figures S7c and S7d) are both the maximum values for each gene across clusters according to equations 1 and 2 in the main text.

For binary classification comparisons on the data from *Velasco et al.*[17], *Islam et al.*[21] and *Zheng et al.*[14], we set pos.only to TRUE and logfc.threshold to 0 in order to obtain all differentially up-regulated genes without any further pre-filtering. For the purpose of this comparison, we also thresholded genesorteR's getPValues output to genes with positive log-fold change values using the same average log fold change formula used by Seurat[26]. We used an adjusted p-value of 0.05 to obtain differentially expressed genes for all methods.

#### - K562 and HEK293T Single Cell ATAC-Seq Data

**Data Generation.** K562 and HEK293T were obtained from ATCC (CCL243 and CRL3216 respectively) and frozen in complete medium supplemented with 10%DMSO at passage 3 and 2 respectively. Cells were thawed and cultured in RPMI1650 medium (Gibco, #31870-025) supplemented with 10% fetal bovine serum and DMEM+Glutamax medium (Gibco, #31966-021) supplemented with 10% fetal bovine serum for K562 and HEK293T respectively. Cells were collected at log growth phase and 1 million cells (300000 HEK293T and 700000 K562) were resuspended in PBS and filtered through Flowmi strainers (40um, Sigma BAH136800040) to remove cell debris. Nuclei were isolated according to the Omni-ATAC lysis buffer formula[27]. Isolated nuclei were resuspended in 10X Genomics nuclei buffer (10X Genomics PN 2000153) and the 10x scATAC-Seq user guide was followed using 10x ATAC-Seq v1.0 kit to produce a single cell ATAC-Seq dataset targeting 6000 nuclei. 10 PCR cycles were applied to the library at the exponential PCR step.

**Data Processing.** Fastq files were processed using Cellranger ATAC version 1 (10X Genomics). The resulting "possorted.bam" file was then filtered for concordantly paired read pairs and processed through the snaptools python module version 1.4.7[28] to produce read counts at different bin resolutions in single nuclei setting --max-num to 10000, –min-cov to 1000, --max-flen to 1000 and –min-flen to 50. To filter nuclei, genomic bin nuclei read count matrices for 10000 candidate nuclei across 617669 genomic bins (5k sized bins) were imported into R using the SnapATAC R package version 1.0.0[28]. We followed the following filtering steps to select confident cells and informative genomic bins. First, log10 total read counts for nuclei were clustered into two clusters using mclust R package version setting modelNames to "E"[2]. Cells that belonged to the cluster with the higher counts was kept. We then removed bins that had less than 1 read in 0.5% of nuclei on average. We then filtered nuclei based on the percentage of reads occurring in the top 500 features. Nuclei whose percentage of reads occurring in the top 500 bins was higher than 4 times the absolute median deviation of all nuclei were removed. Finally, we propose that the total number of reads in nuclei should correlate with the total number of detected bins. Nuclei that did not follow this relationship were filtered by clustering nuclei using a bivariate Gaussian mixture model on log10 total counts and log10 total detected bins using mclust R package version 5.4.3[2] setting modelNames to "VEE". We filtered out the nuclei belonging to the cluster with the lower number of nuclei. This left 529351 genomic bins across 6186 nuclei with an average of 12559 bins per nucleus. See Supplementary Figure S15.

**Semi-Supervised Clustering.** We expected our data to contain primarily two clusters (HEK293T and K562 at roughly 3:7 ratio) as well as potentially ambiguous nuclei resulting from doublets or nuclei debris. To assign nuclei in a semi-supervised manner we used cell line-specific open chromatin regions obtained from HEK293T and K562 DNase-Seq data from the ENCODE consortium[29]. Fastq files were obtained from the UCSC ENCODE website (http://www.genome.ucsc.edu/ENCODE/).

DNase-Seq reads were mapped to the hg38 genome using bowtie2 version 2.2.6[20] and reads that did not align uniquely or had more than 2 mismatches were filtered out. Potential PCR duplicates were removed using samtools rmdup version 1.7[30]. All replicates from the same cell line were concatenated together and peaks were called using JAMM[31] version 1.0.7.5 setting -f 1 -b 100 and -e auto. Peaks from the all.peaks.narrowPeak list from both cell lines that were at least 50bp in width were merged into a single BED file using bedtools merge version 2.25.0[32] and a peak / sample count matrix was obtained for each replicate using bedtools multicov version 2.25.0[32]. We then filtered out peaks that had less than 5 reads per sample on average and used the remaining peaks to perform TMM count normalization and differential expression analysis using the edgeR R package version 3.24.3[16]. Peaks that were differentially expressed with a p-value smaller than 0.001 and absolute average fold-change value bigger than 1 were considered differentially expressed. This resulted in 36531 K562-specific peaks and 39168 HEK293-specific peaks.

To assign single nuclei to either K562 or HEK293T, we intersected the bins detected in each nucleus with both K562-specific peaks and HEK293T-specific peaks. This resulted in a matrix of K562-specific and HEK293T-specific peak intersection counts across all nuclei. Running Singular Value Decomposition on this matrix shows that the first left singular vector separates two clear nuclei groups and a minority of "in between" nuclei. We used the k-means algorithm to cluster the nuclei along the first left singular vector into 3 group (see Main Figure 4). This resulted in 3959 K562 nuclei (64%), 1883 HEK293T nuclei (30.44%) and 344 ambiguous nuclei (5.56%).

We only considered non-ambiguous nuclei for genesorteR and SnapATAC[28] accuracy analysis and used the same DNase-Seq peak sets (see above) as ground truth. We ran genesorteR in all such cases setting binarizeMethod to "naïve" and numPerm for getPValues to 20.

#### - Internet Movie DataBase (IMDB)

IMDB Pajek data (www.imdb.com)[33] were obtained from the SuiteSparse Matrix Collection at Texas A&M University (https://sparse.tamu.edu/Pajek/IMDB). Titles belonging to unknown or unassigned genres, war films and documentaries were removed. sortGenes() was run on the resulting matrix setting binarizeMethod to naive.
